# Suppressing PDGFRβ Signaling Enhances Myocyte Fusion to Promote Skeletal Muscle Regeneration

**DOI:** 10.1101/2024.10.15.618247

**Authors:** Siwen Xue, Abigail M. Benvie, Jamie E. Blum, Nikolai J Kolba, Benjamin D. Cosgrove, Anna Thalacker-Mercer, Daniel C. Berry

**Affiliations:** The Divisional of Nutritional Sciences at Cornell University, Ithaca, NY; Meinig School of Biomedical Engineering, Cornell University, Ithaca, NY; Department of Cell, Development and Integrative Biology, University of Alabama at Birmingham, Birmingham, Alabama; Department of Chemical Engineering; Stanford University; Stanford, CA

## Abstract

Muscle cell fusion is critical for forming and maintaining multinucleated myotubes during skeletal muscle development and regeneration. However, the molecular mechanisms directing cell-cell fusion are not fully understood. Here, we identify platelet-derived growth factor receptor beta (PDGFRβ) signaling as a key modulator of myocyte fusion in adult muscle cells. Our findings demonstrate that genetic deletion of *Pdgfrβ* enhances muscle regeneration and increases myofiber size, whereas PDGFRβ activation impairs muscle repair. Inhibition of PDGFRβ activity promotes myonuclear accretion in both mouse and human myotubes, whereas PDGFRβ activation stalls myotube development by preventing cell spreading to limit fusion potential. Transcriptomics analysis show that PDGFRβ signaling cooperates with TGFβ signaling to direct myocyte size and fusion. Mechanistically, PDGFRβ signaling requires STAT1 activation, and blocking STAT1 phosphorylation enhances myofiber repair and size during regeneration. Collectively, PDGFRβ signaling acts as a regenerative checkpoint and represents a potential clinical target to rapidly boost skeletal muscle repair.

## INTRODUCTION

Adult skeletal muscle contains numerous myofibers supporting posture, movement, and metabolism. The development and maintenance of these fibers rely on the fusion of mononucleated muscle stem cells (MuSCs) or satellite cells^1,2^. MuSCs reside in a quiescent state surrounding the myofibers but, upon stress or injury, can become activated^3–7^. Activated MuSCs can self-renew or become transamplifying myoblasts, which differentiate into fusion-competent myocytes. Myocytes can then fuse with other myocytes or existing myofibers, effectively repairing the injured myofibers^1,3–9^.

Cell fusion is an essential multistep process involving cell cycle exit, migration, and cell-cell interaction that occurs in various biological processes across several tissues and organisms^10–12^. Additionally, fusogenic cells undergo significant plasma membrane remodeling and actin-cytoskeleton reorganization that create fusion protrusions and synapses that allow cell merging^10–18^. Although myocytes express generalized fusion machinery, they also express unique muscle-specific fusogens such as myomaker and myomerger (myomixer/minion)^19–24^. Even though myomaker and myomerger are necessary for myocyte merging, increasing their expression does not always enhance myocyte fusion and regeneration^25^. This highlights the requirement for multiple regulatory and remodeling pathways required for efficient fusion. In agreement with this notion, transforming growth factor beta (TGFβ) signaling prevents fusion by modulating WNT/β-catenin signaling pathways and numerous actin-cytoskeleton regulatory genes^26,27^. Nevertheless, the full spectrum of signaling pathways facilitating myocyte membrane and actin-cytoskeleton remodeling to coordinate fusion remains largely unidentified.

Platelet-derived growth factor receptor beta (PDGFRβ), a receptor tyrosine kinase, has been shown to regulate cell proliferation, migration, and differentiation within various tissues^28–31^. Canonical PDGFRβ signaling within pericytes and smooth muscle cells controls vascular integrity and promotes blood vessel formation and expansion^28,32^. In skeletal muscle, pericytes are critical for skeletal muscle development by promoting myocyte differentiation and limiting MuSC quiescence^33^. More specifically, in vitro, PDGF ligands can stimulate myoblast amplification with inconsistent observations on myotube development^34–36^. Yet, lineage marked *Pdgfrβ*+ cells can contribute to myofiber development and repair^37^. Interestingly, a rare genetic disease, Kosaki overgrowth syndrome, results from single point mutations in the *Pdgfrβ* gene leading to constitutive activation of PDGFRβ signaling accelerating skeletal growth^38–40^. Additionally, patients can develop scoliosis and infantile myofibromas, suggesting possible muscle defects^38–40^. However, muscle-specific genetic necessity and sufficiency tests on PDGFRβ function and signaling on muscle cell proliferation, differentiation, and fusion are lacking.

In this study, we sought to determine the role of PDGFRβ signaling in skeletal muscle biology by employing muscle-specific *Pdgfrβ* genetic necessity and sufficiency tests along with PDGFRβ pharmacological approaches during muscle cell activation, differentiation, and regeneration. We show that the activation of PDGFRβ signaling blunts myotube formation and myofiber regeneration, independent of muscle cell renewal or differentiation. Deleting *Pdgfrβ* in myocytes promotes myotube formation, myonuclear accretion, and fusion, whereas constitutively activating *Pdgfrβ* blunts myofiber regeneration. PDGFRβ signaling utilizes STAT1 phosphorylation to alter TGFβ and focal adhesion gene regulatory networks to mediate myocyte size and fusion. Moreover, pharmacological approaches to inhibit PDGFRβ signaling in murine and human models demonstrate the possibility of targeting PDGFRβ activation to rapidly augment skeletal muscle regeneration.

## RESULTS

### PDGFRβ signaling is active in muscle progenitor cells

To examine the expression pattern of *Pdgfrβ* in muscle progenitor cells, we FACS isolated quiescent PAX7+ cells from hindlimb (quadriceps, gastrocnemius-soleus, and tibialis anterior) muscle groups. To isolate PAX7+ cells, we employed the tamoxifen-inducible *Pax7-Cre^ERT^*^2^ mouse model combined with the indelible genetic lineage tracing tool, *Rosa26^tdTomato^* (Pax7^tdTomato^) (Supplemental Figure 1A)^41,42^. To induce muscle progenitor tdTomato labeling, we administered one dose of tamoxifen (TMX) for two successive days. Notably, we observed high correspondence between reporter and endogenous PAX7 expression, suggesting faithful labeling between endogenous expression and the genetic tool (Supplemental Figure 1B-G). Mice were then randomized into uninjured (quiescent state) or chemically induced (1.2% BaCl_2_) injury groups. Next, we FACS isolated Pax7^tdTomato^ positive and negative cells at 0- and three-days post injury (d.p.i.) to analyze muscle progenitors for *Pdgfrβ* mRNA expression (Figure 1A). *Pdgfrβ* expression was barely detectable within the quiescent Pax7-positive cells compared to the surrounding muscle stroma (Figure 1B). At three d.p.i., a time when MuSCs are activated^1^, we found that *Pdgfrβ* mRNA expression increased within Pax7-positive cells, suggesting a role in the muscle progenitor activated state (Figure 1C). To confirm our RNA expression analysis, we used FACS based approaches to evaluate PDGFRβ protein expression on Pax7^tdTomato^ cells. Indeed, activated Pax7-postive cells (three d.p.i.) stained PDGFRβ positive compared to quiescent Pax7-positive cells, suggesting that PDGFRβ expression is responsive to muscle stress (Figure 1D).

**Figure 1.**
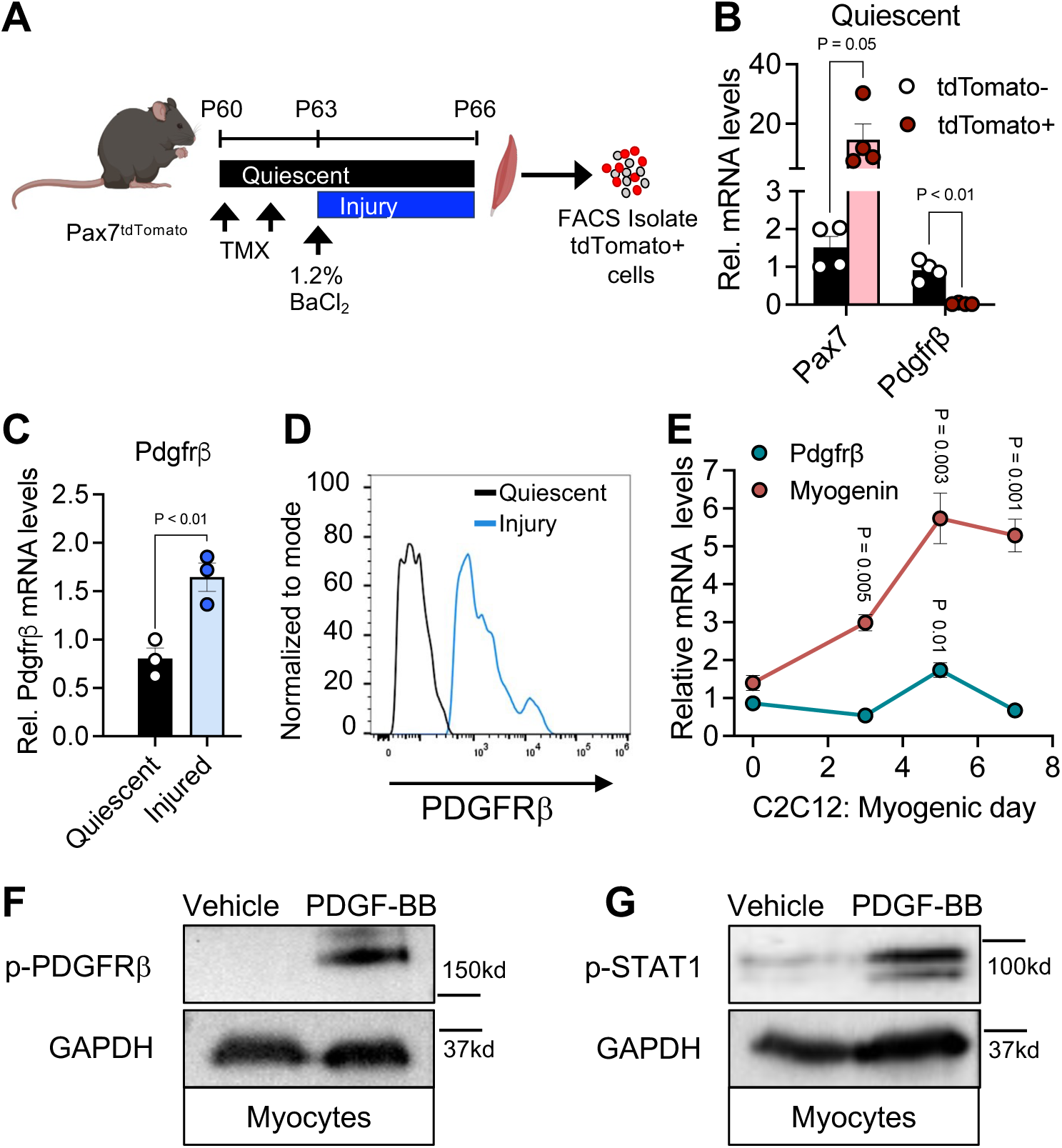
Pdgfrβ is expressed and activated in stimulated muscle progenitor cells. (A) Schematic of the experimental approach. Pax7^tdTomato^ male mice were administered tamoxifen (TMX) at postnatal day 60 (P60) to induce Cre-mediated recombination. Following TMX administration, mice were either uninjured or subjected to a single intramuscular injection of 1.2% BaCl₂ to induce injury in the TA muscle. The TA muscles were harvested, and muscle cells were isolated. Fluorescence-activated cell sorting (FACS) was then employed to separate Pax7^tdTomato^-positive and -negative cells. (B) Quantitative PCR (qPCR) analysis of *Pax7* and *Pdgfrβ* mRNA expression in quiescent Pax7^tdTomato^-positive and -negative cells isolated from the mice described in (A), providing insights into gene expression patterns in muscle progenitor cells (n = 4 biologically independent mice/group). (C) Comparative mRNA expression analysis of *Pdgfrβ* within Pax7^tdTomato+^ cells under quiescent and injured conditions, highlighting the dynamic upregulation of *Pdgfrβ* following muscle injury in the mice described in (A) (n = 3 biologically independent mice/group). (D) Representative flow cytometric histogram showing PDGFRβ expression on PAX7+ cells derived from quiescent and injured TA muscle, as described in (A). (E) Temporal mRNA expression of *Pdgfrβ* and *Myogenin* at specified stages of myogenic differentiation in C2C12 muscle cell line (n = 4 independent cultures per group). (F) Immunoblot analysis of phosphorylated PDGFRβ in myocytes treated with vehicle or PDGF-BB (15 ng/mL for 15 min), demonstrating the activation of PDGFRβ signaling. (G) Immunoblot analysis of phosphorylated STAT1 in myocytes treated with vehicle or PDGF-BB (15 ng/mL for 15 min), indicating downstream signaling effects of PDGFRB activation. Data are presented as mean ± SEM, with each experimental group comprising biological replicates as indicated. Statistical significance was determined using unpaired two-tailed Student’s *t* test (B, C, and E).

To evaluate PDGFRβ’s involvement within the muscle lineage, we first profiled *Pdgfrβ* expression across in vitro myogenesis. We cultured C2C12 cells, a myogenic cell line^43,44^, and then induced them to form myotubes and RNA was harvested at 0-, 3-, 5-, and 7-days post-induction. We found that *Pdgfrβ* expression tracked with *myogenin* but was quickly downregulated upon myotube maturation (Figure 1E). To understand if PDGFRβ signaling could be activated during myogenesis, we isolated primary muscle progenitor cells from Pax7^tdTomato^ control mice. We stimulated them to differentiate into myocytes for four days and subsequently treated them with PDGF-BB, a predominant PDGFRβ activating ligand dimer^29^, for 15 minutes. Immunoblot detection revealed that PDGFRβ could be phosphorylated under these conditions (Figure 1F). The activated PDGFRβ receptor complex is known to activate several downstream effectors such as STAT1^38,45^. Notably, we and others have shown that STAT1 mediates several PDGFRβ-induced biological and metabolic phenotypes that appear independent of cellular proliferation^38,45^. Immunoblotting revealed more STAT1 phosphorylation in response to PDGF-BB ligand addition (Figure 1G). Overall, these data suggest that PDGFRβ is upregulated and stimulated in activated muscle progenitors.

### PDGFRβ activity alters myotube nuclear accretion

To determine whether PDGFRβ has a functional role in myotube development, we examined in vitro myotube formation under conditions of PDGFRβ activation and inhibition. Primary murine muscle progenitor cells were isolated from the hindlimb muscles of male C57BL6/J-129SV mice^46–48^. Cells were cultured, expanded, and then induced to differentiate. Upon differentiation media addition, muscle cells were treated daily for five days with vehicle, PDGF-BB (25 ng/ml), or SU16f (1 µM), a potent and selective inhibitor of PDGFRβ^49^, and myotube formation was assessed (Figure 2A). Activation of PDGFRβ significantly reduced the appearance of myotube development, whereas inhibition of PDGFRβ with SU16f markedly increased the formation of multinucleated myotubes (Figure 2B). Additionally, blocking PDGFRβ signaling increased the fusion index of nucleated myotubes (Figure 2C). Not only did inhibiting PDGFRβ produce larger myotubes but also increased myotube length and diameter (Figure 2D and Supplemental Figure 2A and 2B). In contrast, activating PDGFRβ signaling significantly reduced fusion and predominantly yielded nascent myotubes with 1-2 nuclei per tube (Figure 2C, 2D and Supplemental Figure 2A and 2B). A potential confounder of impacting PDGFRβ signaling could be altered myoblast differentiation, unrelated to muscle cell fusion. To assess this notion, we employed the *Myogenin*-*Cre* (MyoG) mouse model combined with the *Rosa26^tdTomato^* model for myocyte labeling, providing ease for visualizing myocytes and scoring myotube formation (Supplemental Figure 2C-E)^50^. We isolated muscle progenitors from hindlimb muscle groups from MyoG^tdTomato^ mice. Notably, isolation of primary muscle cells from *MyoG^tdTomato^* mice were tdTomato-negative, suggesting that *Myogenin* is not actively expressed in freshly isolated muscle progenitor cells (Supplemental Figure 2D)^51^. Conversely, when differentiated and fused, tdTomato is easily observed in myotubes and corresponds with the mature myotube marker, myosin heavy chain (MyHC) (Supplemental Figure 2E)^52^. Isolated muscle progenitors were cultured at low density and induced with ligands along with differentiation media^26^. We found that neither activating nor inhibiting PDGFRβ affected the formation of MyoG+ myocytes, suggesting that PDGFRβ might not regulate myocyte differentiation but rather myogenic fusion (Figure 2E and Supplemental Figure 2F and 2G).

**Figure 2.**
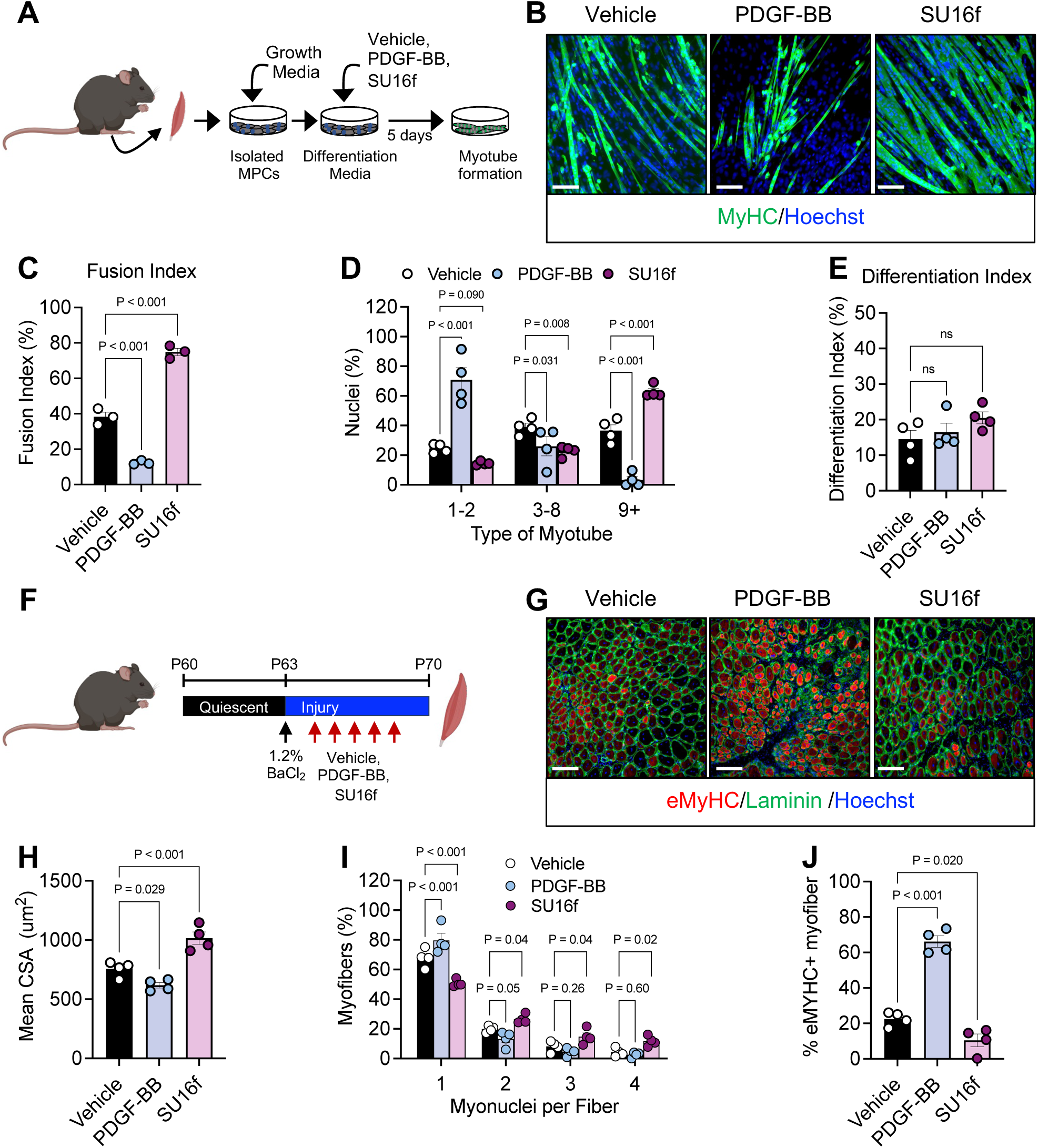
Pdgfrβ activity regulates myotube development and muscle regeneration. (A) Schematic overview of the experimental procedure. Hindlimb muscles were collected from P30 male mice, and isolated muscle progenitor cells (MPCs) were cultured. Throughout differentiation, cells were treated with vehicle, PDGF-BB (25 ng/mL), or SU16f (1 µM) for five days and then myotube formation was assessed. (B) Representative images of myosin heavy chain (MyHC) showing myotube development in cultures as described in (A), providing visual evidence of the effects of PDGF-BB and SU16f on myotube formation. (C) Quantification of the fusion index of differentiated cultures described in (B), reflecting the efficiency of myoblast fusion into myotubes in the treated cultures (n = 3 biologically independent mice per group). (D) Quantification of myotube nuclei count and distribution from cultures described in (B) (n = 4 biologically independent mice per group). (E) Muscle progenitor cells were isolated from hindlimb muscles of MyoG^tdTomato^ mice and cultured at low density. Cells were treated with vehicle, PDGF-BB (25 ng/mL), or SU16f (1 µM), and the differentiation index was determined by monitoring MyoG^tdTomato^ positivity (n = 4 biologically independent mice per group). (F) Schematic overview of the regenerative experimental procedure. MyoG^tdTomato^ mice were subjected to a single intramuscular injection of 1.2% BaCl₂ to induce injury in the TA muscle. Subsequently, mice were administered one dose of vehicle, PDGF-BB (50 ng/mouse), or SU16f (2 mg/Kg) for five consecutive days by intraperitoneal injection. Muscle regeneration was assessed at seven d.p.i. (G) Representative images of TA muscle sections stained for eMyHC and laminin, showing the extent of muscle regeneration and myofiber repair in the different experimental conditions described in (F). (H) Quantification of injured myofiber cross sectional area (CSA) from the images described in (G), providing a comparative analysis of muscle regeneration efficiency (n = 4 biologically independent mice per group). (I) Quantification of myonuclear accretion per injured myofiber, offering insights into the cellular response to injury and subsequent repair (n = 4 biologically independent mice per group). (J) Quantification of eMyHC immunostaining from muscle sections described in (G), indicating the level of ongoing myofiber regeneration in the different experimental conditions (n = 4 biologically independent mice per group). Data are presented as mean ± SEM, with each experimental group consisting of biologically independent replicates as indicated. Statistical significance was determined using a one-way ANOVA for panels (C, E, H, and J) or a two-way ANOVA for panels (D) and (I). Scale bar = 100 µm.

Consistent with our findings in primary muscle cell cultures, treatment of C2C12 cells with PDGF-BB markedly inhibited myotube formation (Supplemental Figure 2H-J). Conversely, blocking PDGFRβ signaling significantly increased the number of multinucleated myotubes (Supplemental Figure 2H-J). To further investigate the role of PDGFRβ in myotube formation, we conducted *Pdgfrβ* shRNA knockdown experiments in C2C12 cells. Like chemical inhibition, *Pdgfrβ* knockdown led to a substantial increase in the number of nuclei within MyHC-positive myotubes (Supplemental Figure 2K, and 2L). Overall, PDGFRβ activation appears to impede myotube formation, while inhibition of PDGFRβ promotes myotube development and enhances myonuclear accretion.

### Inhibiting PDGFRβ promotes myofiber regeneration

The dynamic outcome of activating or inhibiting PDGFRβ on myotube formation prompted us to investigate in vivo targeting of PDGFRβ activity on myofiber regeneration. Towards this end, we performed chemical injury on TA muscle groups by intramuscularly injecting 1.2% BaCl_2_. Simultaneously, we administered PDGF-BB (50 ng/mouse) or SU16f (2 mg/Kg) by intraperitoneal injection for five days and assessed muscle regeneration at seven d.p.i. (Figure 2F). We found that SU16f treatment increased the cross-sectional area (CSA) of injured myofibers (Figure 2G and 2H). In agreement with larger CSA, we observed that SU16f treatment increased myofiber nuclear accretion within injured TA fibers, suggesting the possibility of more fusion events (Figure 2G and 2I). We also evaluated if SU16f augmented myofiber regeneration by staining muscle sections for embryonic myosin heavy chain (eMyHC), a marker of non-regenerated muscles^52,53^. We found that inhibiting PDGFRβ reduced the number of eMyHC myofibers, suggesting enhanced myofiber repair after injury (Figure 2G and 2J). In contrast, PDGF-BB treatment strikingly changed myofiber organization and size (Figure 2G and 2H). PDGF-BB treatment produced smaller regenerating myofibers, characterized by centralized nuclei, and was associated with less myofiber nuclear accretion (Figure 2G and 2I). Changes in myofiber size and nuclear number also were associated with more eMyHC+ myofibers from mice treated with the PDGFRβ ligand, suggesting a delay in repair (Figure 2G and 2J). Together, these data indicate that blocking PDGFRβ activity boosts myotube formation and myofiber regeneration whereas, activating PDGFRβ blunts myotube development and slows myofiber repair.

### Modulating *Pdgfrβ* expression alters skeletal muscle regeneration

Because pharmacological administration of PDGFRβ activators and inhibitors can have systemic off-target effects, we choose to investigate the effects of *Pdgfrβ* gene ablation in muscle cells during regeneration. To do so, we employed a genetic approach by crossing *Pdgfrβ* conditionally floxed (*Pdgfrβ^fl/fl^*)^30^ mice with the *Pax7-Cre^ERT^*^2^ mouse model (Pdgfrβ^Pax7^KO). Additionally, to trace Pax7-positive cells and their progeny, we incorporated the *Rosa26^tdTomato^* lineage reporter into this mouse model (Supplemental Figure S3A). Genetic recombination was induced by administering TMX for two consecutive days. Control mice, carrying the relevant alleles, were also treated with TMX to enable fate mapping analysis and muscle progenitor analyses. Male mice were then randomized into two groups: uninjured and chemically induced muscle injury (1.2% BaCl_2_) and myofibers were assessed seven days later (Figure 3A). In the resting state, myofiber size appeared comparable between control and mutant groups (Supplemental Figure S3B-D). However, following injury, we observed that *Pdgfrβ* deletion increased the CSA of regenerating myofibers (Figure 3B and 3C). Additionally, deleting *Pdgfrβ* within muscle progenitor cells increased myofiber nuclear accretion within regenerating myofibers (Figure 3B and 3D). Consistently, we observed a lack of eMyHC immunostaining within Pdgfrβ^Pax7^KO muscle sections, indicating that the loss of *Pdgfrβ* enhanced myofiber regeneration (Figure 3B and 3E). Of note, we observed similar trends in Pdgfrβ^Pax7^KO female mice after injury (Supplemental Fig. 3E-G).

**Figure 3.**
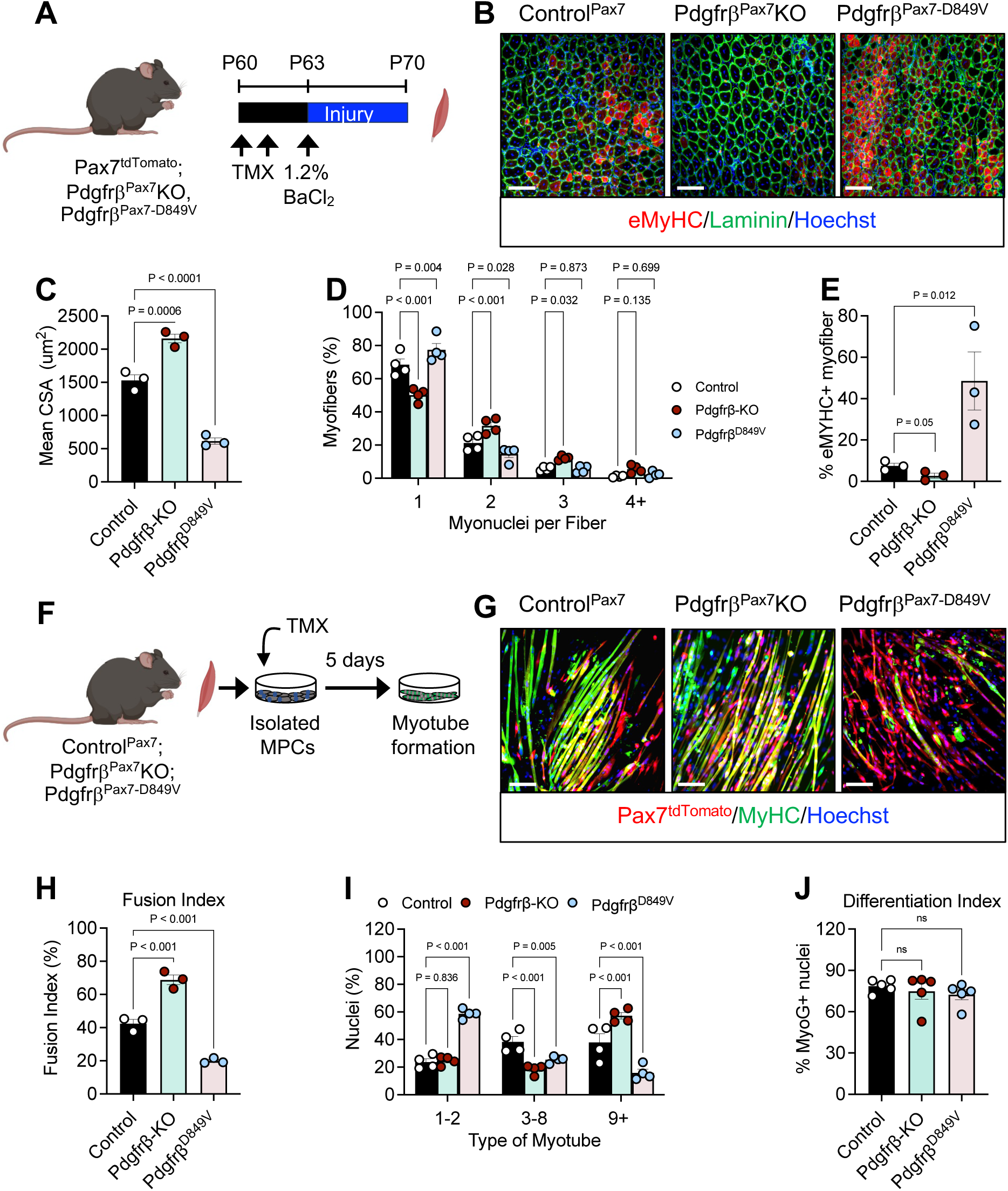
Genetically altering *Pdgfrβ* expression changes muscle regeneration and myotube development. (A) Schematic of the experimental approach. Control^Pax^^7^, Pdgfrβ^Pax7^KO, and Pdgfrβ^Pax7-D849V^ mice were administered TMX at postnatal day 60 (P60) to induce Cre-mediated recombination. TA muscles were subsequently injured with 1.2% BaCl₂ and analyzed at seven d.p.i. (B) Representative images of injured TA muscle sections immunostained for eMyHC and laminin from mice described in (A), illustrating the regenerative response across different genotypes. (C) Quantification of mean cross-sectional area (CSA) of injured TA myofibers from sections described in (B) from mice described in (A), reflecting the differences in muscle regeneration (n = 3 biologically independent mice per group). (D) Quantification of myonuclear accretion within injured myofibers, providing insights into cellular fusion and muscle repair dynamics in the different experimental groups (n = 4 biologically independent mice per group). (E) Quantification of eMyHC immunostaining from TA muscle sections described in (B), indicating the level of ongoing regeneration post-injury (n = 3 biologically independent mice per group). (F) Schematic overview of the in vitro experimental approach. Muscle progenitor cells were isolated from the hindlimb muscle groups of Control^Pax^^7^, Pdgfrβ^Pax7^KO, and Pdgfrβ^Pax7-D849V^ male mice. After isolation, cells were administered TMX to induce recombination, expanded, and subsequently differentiated. Myotube development was assessed in culture. (G) Representative images showing myotube development as assessed by tdTomato (Pax7) fluorescence and MyHC staining from cultures derived from mice described in (F). (H) Quantification of the fusion index from the images described in (G) and cultures described in (F), reflecting the efficiency of muscle cell fusion into myotubes (n = 3 biologically independent mice per group). (I) Quantification of myotube nuclei count and distribution from the images described in (G) from cultures described in (F) (n = 4 biologically independent mice per group). (J) Quantification of the differentiation index from low density muscle cell cultures. Muscle progenitors were isolated from hindlimb muscle groups from of Control^Pax^^7^, Pdgfrβ^Pax7^KO, and Pdgfrβ^Pax7-D849V^. After isolation, low density cultures were administered TMX to induce recombination, expanded, and subsequently differentiated. Colocalization of tdTomato fluorescence and myogenin-positive cells were used to calculate the differentiation index (n = 5 biologically independent mice per group). Data are presented as mean ± SEM, with each experimental group consisting of biologically independent replicates as indicated. Statistical significance was determined using a one-way ANOVA for panels (C, E, H, and J) or a two-way ANOVA for panels (D) and (I). Scale bar = 100 µm.

To investigate whether enhanced basal PDGFRβ signaling negatively affects myofiber regeneration, we employed an inducible constitutively activated *Pdgfrβ* gain-of-function model in muscle cells (Pdgfrβ^Pax7-D849V^)^54^ (Supplemental Figure 3A). This knock-in allele incorporates an activating point mutation within the *Pdgfrβ* kinase domain, resulting in increased basal PDGFRβ signaling independent of ligand presence^54^. Activation of *Pdgfrβ^-D^*^849^*^V^* was achieved by administering TMX to adult mice for two consecutive days. Control and PDGFRβ-activated mice were then randomized into non-injury or chemically induced muscle injury groups (Figure 3A). Under quiescent conditions, TA myofiber size did not differ between groups (Supplemental Fig. S3B-D). However, following injury (seven d.p.i.), muscle sections from Pdgfrβ^Pax7-D849V^ mice displayed distorted tissue architecture and the presence of micromyofibers with significantly smaller myofiber CSAs (Figure 3B and 3C). This impairment in myofiber regeneration was further evident by a reduced number of nuclei per regenerating myofiber in Pdgfrβ^Pax7-D849V^ samples, suggesting a delay in muscle repair response (Figure 3B and 3D). In agreement with this notion, eMyHC immunostaining was markedly elevated in Pdgfrβ^Pax7-D849V^ sections compared to control samples (Figure 3B and 3E). Notably, female Pdgfrβ^Pax7-D849V^ injured muscle sections revealed similar trends (Supplemental Fig. 3E-G). These data collectively suggest that deletion of *Pdgfrβ* in muscle progenitor cells facilitates myofiber repair, whereas activation of *Pdgfrβ* hinders myofiber regeneration.

### Modulating *Pdgfrβ* expression alters in vitro myotube development

To continue to evaluate the role in PDGFRβ in myotube development, we isolated muscle progenitors from hindlimb muscle groups from control, Pdgfrβ^Pax7^KO, and Pdgfrβ^Pax7-D849V^ mice. Upon culturing, we administered TMX to induce recombination and tdTomato labeling and allowed myotubes to form (Figure 3F). We found that deleting *Pdgfrβ* increased the fusion index of nuclei within myotubes (Figure 3G and 3H). Additionally, *Pdgfrβ* deletion was also associated with an increase in the number of nuclei per myotube, indicative of enhanced myonuclear accretion^25^ (Figure 3G and 3I). In contrast, constitutive activation of *Pdgfrβ* appeared to blunt myotube development reducing the fusion index (Figure 3G and 3H). This was further evident based on imaging as many mono-and di-nucleated Pax7^tdTomato^ cells could be observed but did not co-stain with myosin (Figure 3G). Consistently, myonuclear accretion quantification revealed that many Pdgfrβ^Pax7-D849V^ cells remained in the nascent myotube category and few Pdgfrβ^Pax7-D849V^ myotubes contained 9 or more nuclei (Figure 3G and 3I). Overall, the data indicate that muscle progenitor deletion of *Pdgfrβ* results in myonuclear accretion and myotube development whereas, activating *Pdgfrβ* reduces myotube formation and potential fusion.

### PDGFRβ activity does not affect muscle progenitor cell proliferation

To examine how PDGFRβ might regulate in vitro myotube development and in vivo muscle regeneration, we assessed if PDGFRβ signaling altered PAX7 cell numbers. Towards this end, we isolated Pax7^tdTomato+^ cells from TMX-induced control, Pdgfrβ^Pax7^KO and Pdgfrβ^Pax7-D849V^ TA muscle groups at quiescence, and three and seven days after injury. By flow cytometric analysis, we found that neither *Pdgfrβ* deletion nor activation affected Pax7^tdTomato^ numbers at rest or after injury (Supplemental Fig. 3H)^46^. To validate these findings, we performed PAX7 immunostaining of muscle sections from control and Pdgfrβ mutant mice in the injured states (seven d.p.i.). Imaging and quantification revealed comparable PAX7 staining and numbers regardless of genotype (Supplemental Fig. 3I and 3J).

Because we did not observe changes in PAX7 cell numbers, we evaluated if Pdgfrβ activation or deletion affected myocyte differentiation. To test this, we isolated muscle progenitors from control, Pdgfrβ^Pax7^KO, and Pdgfrβ^Pax7-D849V^ hindlimb muscle groups and plated cells at low density^55^. Cells were then administered TMX and differentiated (Supplemental Fig. 3K). Regardless of genotype, we found comparable differentiation index as assessed by MyoG-positivity within Pax7^tdTomato^ cells (Figure 3J and Supplemental Figure 3L). Together, it appears that Pdgfrβ activity does not directly impact muscle progenitor number nor myocyte differentiation, suggesting that Pdgfrβ activity may regulate myocyte fusion.

### Myocytes defective in PDGFRβ activity have altered myofiber regenerative capacity

To assess whether PDGFRβ signaling is necessary and sufficient for myocyte fusion during skeletal muscle regeneration, we utilized the MyoG-Cre mouse model in combination with *Pdgfrβ^fl/f^*^l^ mice (Pdgfrβ^MyoG^KO) or *Pdgfrβ^D^*^849^*^V^* (Pdgfrβ^MyoG-D849V^) constitutively activated mice. At postnatal day 60 (P60), we chemically injured the TA muscle group and assessed regeneration at seven d.p.i. (Figure 4A). Histological analyses TA muscle sections revealed that *Pdgfrβ* deletion significantly increased regenerating myofiber size, whereas constitutive activation of *Pdgfrβ* impaired muscle regeneration (Figure 4B and 4C). *Pdgfrβ* deletion also increased the nuclear number in newly repaired myofibers, which was also associated with larger myofiber CSAs (Figure 4B and 4D). Conversely, *Pdgfrβ* activation diminished the number of nuclei per newly formed myofibers with only one to two nuclei per injured myofiber, suggestive of impaired myocyte fusion. Immunostaining for eMyHC indicated enhanced myofiber repair in *Pdgfrβ*-deleted mice compared to controls (Figure 4B and 4E). In contrast, a higher number of eMyHC-positive myofibers were observed in Pdgfrβ^D849V^ mice, suggesting delayed regeneration (Figure 4B and 4E).

**Figure 4.**
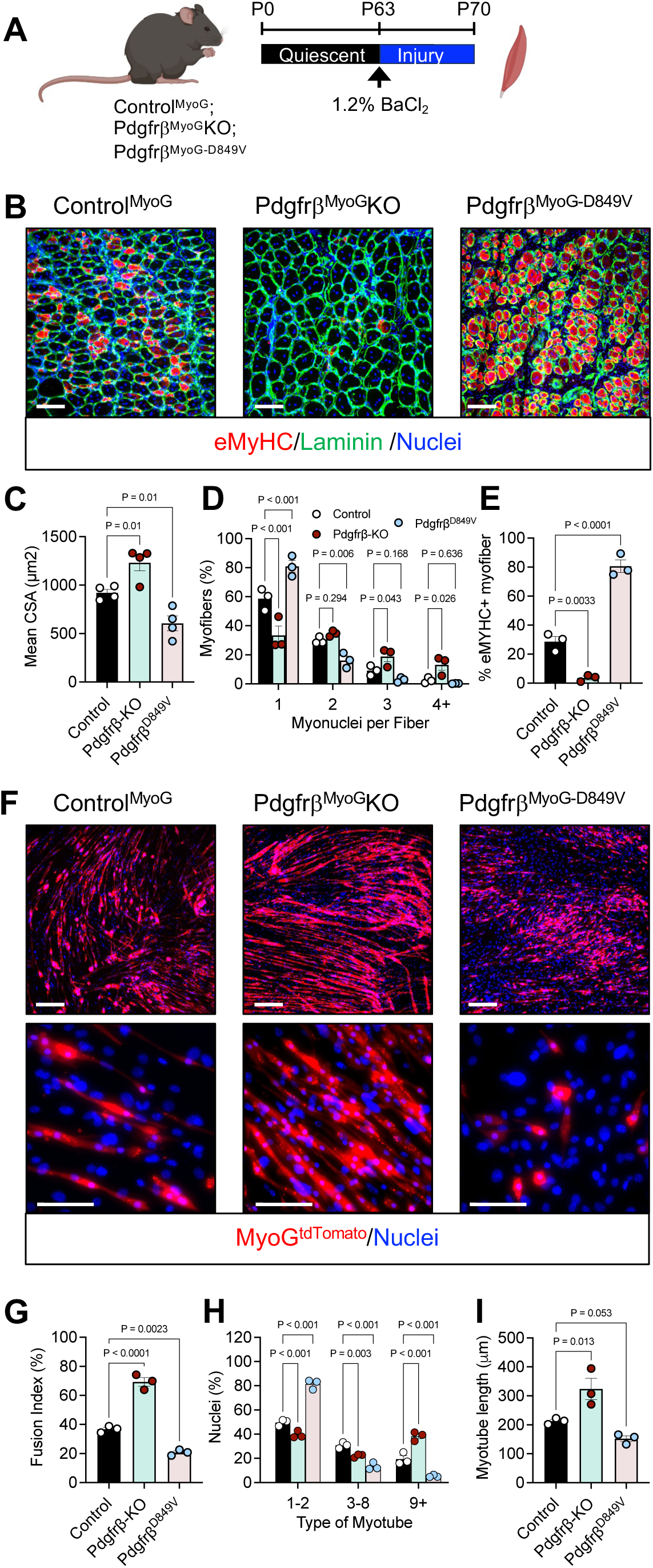
Genetically altering Pdgfrβ activity in myocytes modifies muscle regeneration. (A) Schematic overview of the experimental setup: TA muscles from Control^MyoG^, Pdgfrβ^MyoG^KO, and Pdgfrβ^MyoG-D849V^ mice were intramuscularly injected with 1.2% BaCl₂. Muscle tissues were harvested and analyzed at seven days post-injury (d.p.i.). (B) Representative immunofluorescence images illustrating eMyHC and laminin staining in injured TA muscle sections. The images provide a visual context for the extent of muscle regeneration across the different genotypes. (C) Quantitative assessment of myofiber cross-sectional area (CSA) in the injured regions of TA muscles from sections and images described in (B) (n = 4 biologically independent mice per group). (D) Quantification of myonuclear accretion within injured myofibers, a metric of cellular fusion and muscle repair (n = 3 biologically independent mice per group). (E) Quantification of eMyHC-positive fibers within the injured TA muscle, providing insights into the ongoing myogenic processes in the different experimental groups (n = 3 biologically independent mice per group). (F) Muscle progenitor cells isolated from the hindlimb muscles of Control^MyoG^, Pdgfrβ^MyoG^KO, and Pdgfrβ^MyoG-D849V^ male mice were expanded and differentiated in vitro. Representative images of MyoG^tdTomato^ fluorescence, with higher magnification (40x) presented to detail myotube formation. (G) Fusion index quantification in differentiated cultures, reflecting the efficiency of myoblast fusion into multinucleated myotubes (n = 3 biologically independent mice per group). (H) Quantification of nuclei within myotubes from cultures described in (B) (n = 3 biologically independent mice per group). (I) Measurement of myotube length in differentiated cultures, serving as an indicator of myotube maturation and growth (n = 3 biologically independent mice per group). Data are presented as mean values with individual data points ± S.E.M. Statistical significance was determined using a one-way ANOVA for panels (C-E) and (G-I). Scale bar = 100 µm.

A potential confounder with the MyoG mouse model is that mature myofibers could be affected by *Pdgfrβ* deletion or activation, altering muscle repair. To address this, we generated mice with *Pdgfrβ* deletion (*Pdgfrβ^fl/fl^*) or activation (*D849V*) specifically in mature myofibers using the TMX-inducible *human skeletal actin-MerCreMer* (HSA-MCM) system^56^. At P60, TMX was administered to induce myofiber *Pdgfrβ* deletion or activation, followed by chemical injury to the TA muscle group (Supplemental Figure 4A). Regardless of genotype, regenerating muscles exhibited similar characteristics in terms of myofiber cross-sectional area, eMyHC expression, and myonuclear accretion, suggesting that PDGFRβ signaling in mature myofibers might not significantly affect the regenerative process (Supplemental Figure S4B-D).

Our results showed that mice with the myocyte-specific deletion of *Pdgfrβ* exhibited early signs of repair. Accordingly, we investigated the effect of PDGFRβ activity on the repair of injured myofibers and sampled regenerating muscle at five d.p.i. (Figure S4E)^8^. At this early time point, muscle progenitors are activated, amplifying, differentiating, and begin to fusion. While Pdgfrβ^MyoG^KO five d.p.i. myofibers had comparable CSAs to controls, they contained less eMyHC staining and had more nuclei per injured myofiber (Figure S4F-I). In contrast, regenerating muscles from Pdgfrβ^MyoG-D849V^ mice were disorganized, smaller, and more myofibers were eMyHC-positive, indicating impaired and delayed muscle regeneration (Figure S4F-I). These findings demonstrate that *Pdgfrβ* deletion in myocytes enhances the early stages of muscle repair by promoting myofiber repair and nuclear accretion, whereas constitutive activation of PDGFRβ impairs these processes.

### Fusion defects in adult myocytes with altered PDGFRβ activity

We next assessed if PDGFRβ signaling in myocytes affected the formation of multinucleated myotubes in vitro. To do so, primary muscle cells were isolated from control, Pdgfrβ^MyoG^KO, and Pdgfrβ^MyoG-D849V^ mice and were differentiated for five days and myotubes were assessed. Of note, this model ensures that muscle progenitors remain wild-type for *Pdgfrβ* pending myogenin expression, at the onset of myocyte differentiation (Supplemental Figure 2C-E). Strikingly, we found that Pdgfrβ^MyoG^KO cultures had an increase in the appearance of myotube development, whereas constitutive *Pdgfrβ* activation prevented myotube formation (Figure 4F). Quantification of the fusion index showed more fusion events in the Pdgfrβ^MyoG^KO cultures whereas, Pdgfrβ^MyoG-D849V^ cultures had the least amount of fusion (Figure 4F and 4G). Moreover, nuclei quantification of Pdgfrβ^MyoG^KO cultures revealed a higher number of multinucleated myotubes compared to controls, while Pdgfrβ^MyoG-D849V^ cells remained predominantly mononuclear (Figure 4F-H). In agreement with more nuclear fusion events, Pdgfrβ-KO myotubes were longer, whereas activation of *Pdgfrβ* stunted myotube length (Figure 4I). Taken together, the data suggest that myocyte fusion can be blocked by PDGFRβ signaling.

### Rescue of fusion defects in Pdgfrβ^-D849V^ myocytes

The fusion defects observed in Pdgfrβ^-D849V^ myocytes led us to test if we could rescue Pdgfrβ^MyoG-D849V^ myotube development by performing a cell mixing experiment with wild-type cells^22^. To do so, we co-cultured Pdgfrβ^MyoG-D849V^ muscle cells (tdTomato+) with wild-type cells (no MyoG-Cre or tdTomato) and visualized myotube formation after four days of differentiation (Figure 5A). As expected, control (MyoG^tdTomato^) myocytes fused and formed multinucleated myotubes, whereas Pdgfrβ^MyoG-D849V^ cells in monoculture remained as myocytes with limited myotube development (Figure 5B). Conversely, co-culturing Pdgfrβ^MyoG-D849V^ cells with wild-type cells appeared to rescue the fusion defect induced by constitutive *Pdgfrβ* activation. Imaging revealed that chimeric myotubes displayed a diffuse tdTomato signal throughout, indicating the contribution of Pdgfrβ^MyoG-D849V^ myocytes to myotube formation (Figure 5B). Moreover, co-culturing Pdgfrβ^MyoG-D849V^ cells significantly increased the fusion index (Figure 5C). Increased fusion was also confirmed by nuclei counts, suggesting more multinucleated myotubes and myonuclear accretion (Figure 5D). These results suggest that Pdgfrβ^MyoG-D849V^ myocytes are fusion competent but may be unable to create cellular interactions to cause fusion events.

**Figure 5.**
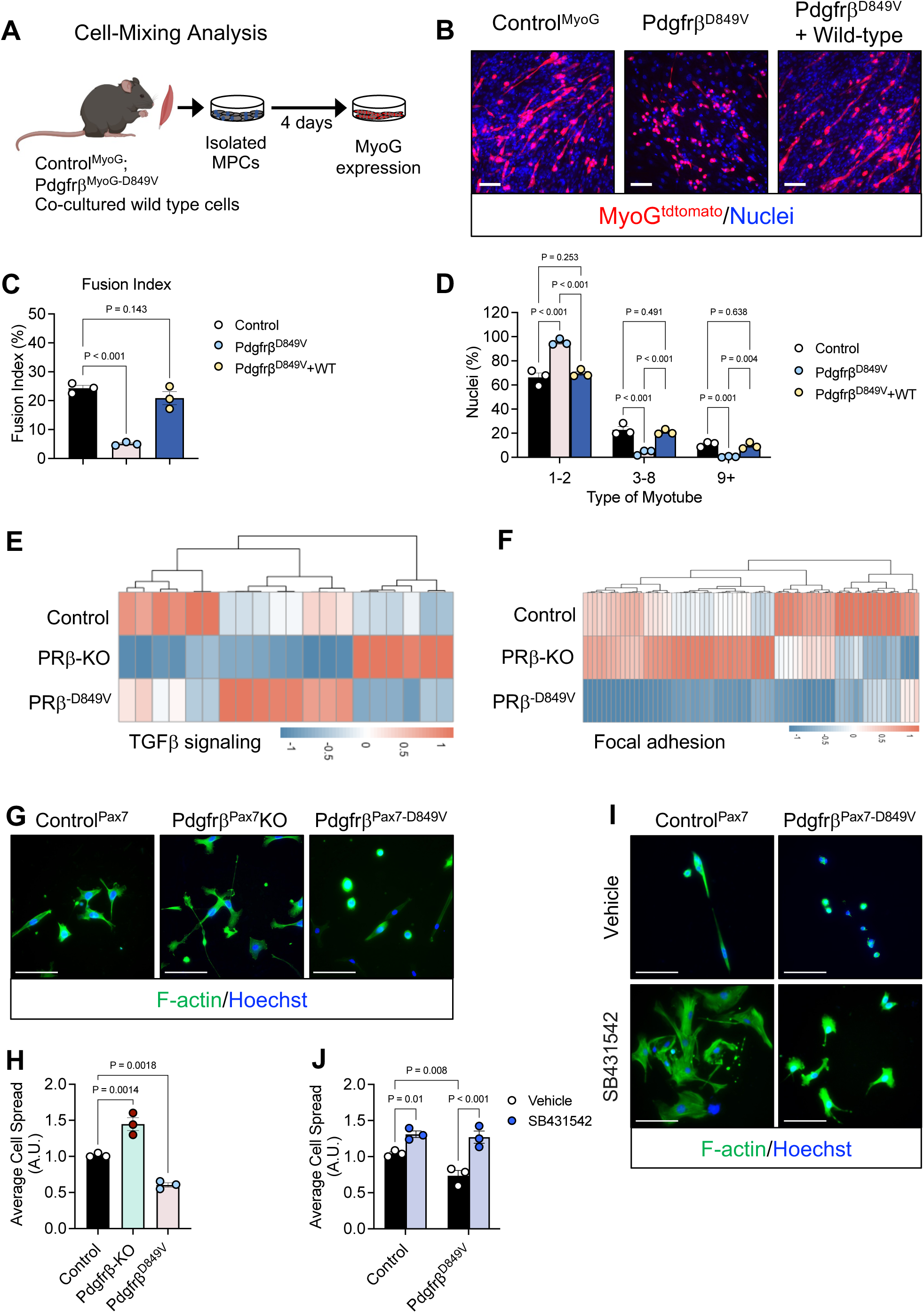
Pdgfrβ signaling cooperates with Tgfβ signaling to control myocyte morphology. (A) Schematic of the in vitro cell mixing experimental procedure. Muscle progenitor cells were isolated from the hindlimb muscle groups of from Control^MyoG^, Pdgfrβ^MyoG-D849V^ and wild-type mice. Control^MyoG^ and Pdgfrβ^MyoG-D849V^ cells were plated grown and differentiated whereas Pdgfrβ^Pax7-D849V^ were co-cultured with wild-type. Myotube and chimeric myotube development was assessed by MyoG^tdTomato^ fluorescence. (B) Representative images of myotube development as assessed by MyoG^tdTomato^ fluorescence from muscle progenitors isolated in (A). (C) Quantification of the fusion index from myotube cultures described in (B), providing a measure of muscle cell fusion in response cell mixing (n = 3 biologically independent mice per group). (D) Quantification of nuclei within myotubes from images described in (B) and cultures described in (A) (n = 3 biologically independent mice per group). (E, F) Heatmap of gene expression profiles comparing hindlimb muscles from Control^MyoG^, Pdgfrβ^MyoG^KO, and Pdgfrβ^MyoG-D849V^ P30 male mice. Gene transcript analysis revealed changes in TGFβ signaling (E) and focal adhesion (F) genes (n = 5 biologically independent mice/group). (G) Representative images of F-actin staining to reveal myocyte morphology from muscle progenitors isolated from hindlimb muscle groups from Control^Pax^^7^, Pdgfrβ^Pax7^KO, and Pdgfrβ^Pax7-D849V^. Cells were grown and differentiated for one day and stained with F-actin. (H) Quantification of cell spreading from myocyte cultures described in (G), providing a measure of muscle cell morphology in response to PDGFRβ activity (n = 3 biologically independent mice per group). (I) Representative images of F-actin staining to reveal myocyte morphology from muscle progenitors isolated from hindlimb muscle groups from Control^Pax^^7^ and Pdgfrβ^Pax7-D849V^. Cells were given TMX, grown, and differentiated for one day in the presence of vehicle or SB431542 (1 µM). Cells were stained with F-actin. (J) Quantification of cell spreading from myocyte cultures described in (I), providing a measure of muscle cell morphology in response to PDGFRβ activity and TGFβ inhibition (n = 3 biologically independent mice per group). Data are presented as mean values with individual data points ± S.E.M. Statistical significance was determined using a one-way ANOVA for panels (C), (D), (H) or a two-way ANOVA for panels (J). Scale bar = 100 µm.

### PDGFRβ signaling alters TGFβ signaling and focal adhesion pathways

To identify gene regulatory networks controlled by PDGFRβ signaling within skeletal muscle, we performed bulk RNA-sequencing from Control^MyoG^, Pdgfrβ^MyoG^KO, and Pdgfrβ^MyoG-D849V^ hindlimb muscle groups in the quiescent state. Analysis of the transcriptomic dataset revealed changes in the relative expression of known TGFβ target genes, which were significantly reduced in the Pdgfrβ^MyoG^KO myocytes but were reciprocally upregulated in the Pdgfrβ^MyoG-D849V^ model (Figure 5E and Supplemental Fig. S5A and S5B). Furthermore, we observed common modulation of gene regulatory networks involved in focal adhesion between Pdgfrβ^MyoG^KO and Pdgfrβ^MyoG-D849V^ myocytes (Figure 5F and Supplemental Fig. S5C and S5D). Interestingly, TGFβ signaling and cytoskeleton remodeling have been shown to block fusion^57^. Notably, actin remodeling has been shown to facilitate the creation of fusion protrusions that allow myocytes to interact, which promotes fusogenic proteins to cooperate and fusion^26,58,59^. To visualize if PDGFRβ activity alters cell morphology and actin alignment, we plated TMX-induced Pdgfrβ^Pax7^KO and Pdgfrβ^Pax7-D849V^ cells at low density and stained them for F-actin one-day post-differentiation^60^. Deleting *Pdgfrβ* promoted cellular spreading for cellular contact and potential fusion points (Figure 5G and 5H). In contrast, activating *Pdgfrβ* blocked cell spreading, where many of the cells remained spherical and did not appear to contact each other (Figure 5G and 5H). To better understand if TGFβ signaling facilitates Pdgfrβ^-D849V^ cell spreading blockade, we plated control and Pdgfrβ^Pax7-D849V^ muscle cells at low density and administered TMX in culture. Cells were then differentiated and treated with vehicle or SB431542 (1 µM), a TGFβ inhibitor^61^, for 24 hours. F-actin staining revealed that blocking TGFβ with SB431542 allowed Pdgfrβ^Pax7-D849V^ cells to spread and create contacts, suggesting a possible cooperation between PDGFRβ and TGFβ signaling pathways (Figure 5I and 5J).

### PDGFRβ-STAT1 signaling regulates myogenic fusion

Next, we wanted to understand what mediates PDGFRβ signaling in muscle cells to prevent cell spreading and fusion. PDGFRβ activation has been shown to be mediated by STAT1, controlling PDGFRβ signaling in beige fat development, fibrosis, inflammation, and hypercholesterolemia^38,45,54,62^. Indeed, the PDGFRβ ligand, PDGF-BB, could induce the phosphorylation of STAT1 in in vitro derived myocytes (Figure 1G). Therefore, we investigated if blocking STAT1 phosphorylation could restore muscle cell fusion in response to PDGFRβ activation in culture. Towards this end, we used the FDA approved drug, fludarabine^63^, used to treat chronic lymphocytic leukemia^64^, to block PDGFRβ-induced STAT1 phosphorylation^45^. While PDGF-BB could stimulate the phosphorylation of PDGFRβ in the presence of fludarabine, STAT1 phosphorylation was blocked, demonstrating that fludarabine interferes with PDGFRβ induced STAT1 phosphorylation (Supplemental Fig. 6A and 6B). Next, we assessed if inhibiting STAT1 phosphorylation could restore myotube formation in PDGF-BB treated C2C12 cells. PDGF-BB treated C2C12s generated myotubes with less nuclei per myotube compared to controls (Supplemental Fig. 6C-E). Notably, fludarabine alone significantly increased the fusion index and myonuclear accretion of C2C12 cells (Supplemental Fig. 6C-E). In contrast, co-treating PDGF-BB cultures with fludarabine produced myotubes containing many more nuclei, which resembled vehicle treated cultures (Supplemental Fig. 6C-E).

Next, we hypothesized that treating Pdgfrβ^D849V^ muscle cell cultures with fludarabine might restore their fusogenic potential. To test this notion, we isolated primary muscle cells from the MyoG mouse models (Control and Pdgfrβ^D849V^) due to the relevance of PDGFRβ in myocyte mediated fusion. Cells were then treated with vehicle or fludarabine and myotube development was assessed (Figure 6A). While the fusion index remained low for Pdgfrβ^D849V^ vehicle treated cultures, fludarabine treatment appeared to rescue myotube formation and increased myotube nuclear accretion (Figure 6B-D). Of note, control cultures treated with fludarabine also had an increase in the number of nuclei per MyHC-positive myotubes (Figure 6D). We confirmed these findings on cultured primary muscle cells from the Pax7-Pdgfrβ^D849V^ gain-of function mouse model (Supplemental Fig. 6F-H). Together, blocking STAT1 restores myotube myonuclear accretion in response to PDGFRβ activation, suggesting that STAT1 may function downstream of PDGFRβ activation.

**Figure 6.**
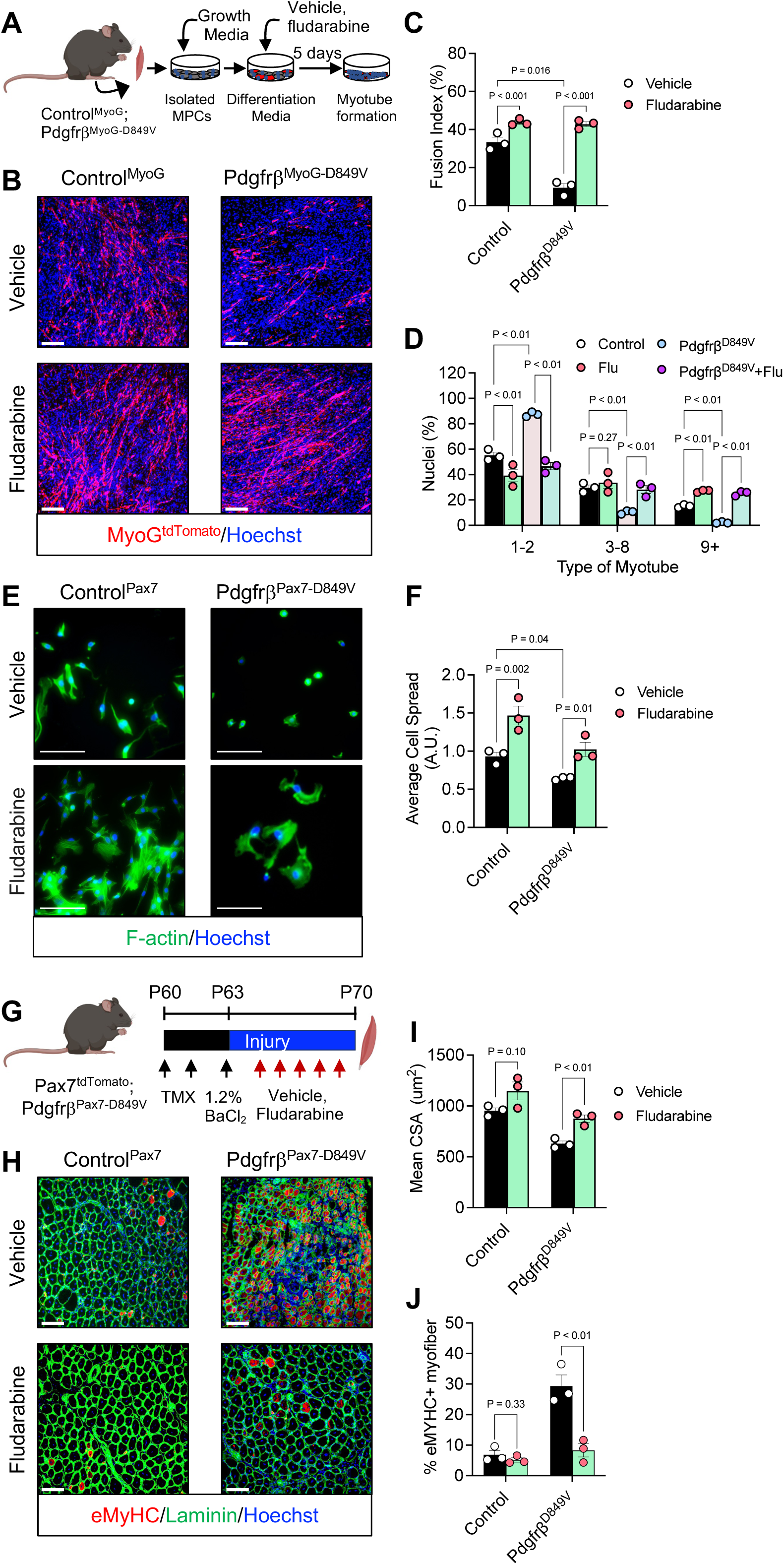
STAT1 mediates PDGFRβ signaling to control myocyte fusion and myofiber regeneration. (A) Schematic of the in vitro experimental procedure. Muscle progenitor cells were isolated from the hindlimb muscle groups from Control^MyoG^ and Pdgfrβ^MyoG-D849V^ male mice. Cells were grown and differentiated in the presence of either vehicle (1% DMSO) or fludarabine (1 µM). Myotube formation was assessed after five days of treatment. (B) Representative images of myotube development as assessed by MyoG^tdTomato^ fluorescence from muscle progenitors isolated in (A). (C) Quantification of the fusion index from myotube cultures described in (B), providing a measure of muscle cell fusion in response to STAT1 inhibition (n = 3 biologically independent mice per group). (D) Quantification of nuclei within myotubes from cultures described in (B) (n = 3 biologically independent mice per group). (E) Muscle progenitor cells were isolated from the hindlimb muscle groups from Control^Pax7^ and Pdgfrβ^Pax7-D849V^ male mice. Cells were grown and differentiated for one day in the presence of vehicle (1% DMSO) or fludarabine (1 µM). Cells were stained with F-actin to visualize cell size and actin orientation. (F) Quantification of cell spreading from myocyte cultures described in (E), providing a measure of muscle cell morphology in response to Stat1 inhibition (n = 3 biologically independent mice per group). (G) Schematic of the in vivo experimental design. Control^Pax^^7^ and Pdgfrβ^Pax7-D849V^ male mice were subjected to TA muscle injury using 1.2% BaCl₂. Post-injury, mice were treated with one dose of vehicle or fludarabine (3 mg/ Kg) for five consecutive days by intraperitoneal injection. Myofiber regeneration was analyzed at seven d.p.i.. (H) Representative images of TA muscle sections from injured mice, immunostained for eMyHC and laminin, showing the extent of muscle regeneration in the different treatment groups as described in (G). (I) Quantification of the cross-sectional area (CSA) of injured TA myofibers from sections described in (H), providing a comparative analysis of muscle regeneration efficiency (n = 3 biologically independent mice per group). (J) Quantification of eMyHC immunostaining from TA muscle sections, indicating the level of ongoing regeneration in the injured muscle as described in (H) (n = 3 biologically independent mice per group). Data are presented as mean values with individual data points ± S.E.M. Statistical significance was determined using a one-way ANOVA for panels (D), (I), and (J) or a two-way ANOVA for panels (C) and (F). Scale bar = 100 µm.

Because fludarabine could restore the fusion potential of constitutively activated Pdgfrβ, we investigated if blocking STAT1 phosphorylation impacted Pdgfrβ^Pax7-D849V^ myocyte spreading and morphology. Towards this end, we plated TMX-induced Control^Pax7^ and Pdgfrβ^Pax7-D849V^ cells at low density and stained them for F-actin one-day post-differentiation. Cells were also treated with vehicle (1% DMSO) or fludarabine (1 µM) during the one-day differentiation period. In control cultures, fludarabine appeared to increase the cell spreading of myocytes (Figure 6E and 6F). Furthermore, treating Pdgfrβ^Pax7-D849V^ cultures with fludarabine was able to restore cell morphology and spreading to control levels, suggesting that STAT1 facilitates *Pdgfrβ*-induce smaller cell morphology to prevent fusion (Figure 6E and 6F).

Our in vitro data suggest that blocking STAT1 signaling can overcome *Pdgfrβ* constitutive activation to promote myocyte spreading and myotube myonuclear accumulation. These data led us to hypothesize that administering fludarabine might improve the regenerative capacity of Pdgfrβ^D849V^ muscles. Towards this end, we performed chemical muscle injury on Control^Pax7^ and Pdgfrβ^Pax7-D849V^ mice; subsequently, mice were randomized to vehicle or fludarabine for five days (Figure 6G). As expected, at seven d.p.i., we found histological disruption of muscle regeneration in Pdgfrβ^Pax7-D849V^ TA muscles (Figure 6H). Moreover, administration of fludarabine appeared to reestablish tissue repair and organization Pdgfrβ^D849V^ mice (Figure 6H). CSA quantification revealed larger Pdgfrβ^Pax7-D849V^ myofibers with centrally localized nuclei after fludarabine treatment (Figure 6I). Myofiber regeneration analysis revealed that eMyHC staining was reduced in fludarabine treated Pdgfrβ^Pax7-D849V^ TA myofibers. Additionally, we observed similar results when we treated Pdgfrβ^MyoG-D849V^ with fludarabine (Supplemental Fig. 6I). These data suggest that STAT1 appears to mediate PDGFRβ myocyte signaling to coordinate myofiber repair (Figure 6H and 6J).

### PDGFRβ signaling regulates human muscle cell fusion

To determine the human application of Pdgfrβ signaling, we evaluated whether pharmacologically manipulating Pdgfrβ influenced human muscle cell fusion and myotube formation. Crude primary human muscle progenitor cells were prepared from isolated vastus lateralis (quadricep) muscle biopsies from young female donors and cultured for four passages^65^. Next, we FACS isolated (CD56+/CD29+) cells to enrich for human muscle progenitors, subsequently cells were plated and passaged twice before experimentation (Figure 7A and Supplemental Fig. 7A-C)^65^. Upon serum restriction, cells were treated with vehicle, PDGF-BB, and SU16f throughout myotube formation. After seven days, we noted that PDGF-BB treatment appeared to block the formation of MyHC+ myotubes (Figure 7B). In agreement, fusion index quantification revealed less myonuclear accretion into MyHC+ myotubes in response to PDGF-BB treatment (Figure 7C). Consistent with reduced fusion, we observed many more nascent myotubes in response to PDGF-BB treatment (Figure 7D). In contrast, treating primary human muscle cells with SU16f increased the fusion index, myotube nuclear accretion, and myotube diameter and length (Figure 7B-F). In agreement with PDGFRβ-STAT1 signaling, we found that blocking STAT1 phosphorylation with fludarabine increased human myotube development and myonuclear accretion (Supplemental Fig. 7D-G). Altogether, these data suggest that blocking PDGFRβ signaling increases human muscle cell fusion.

**Figure 7.**
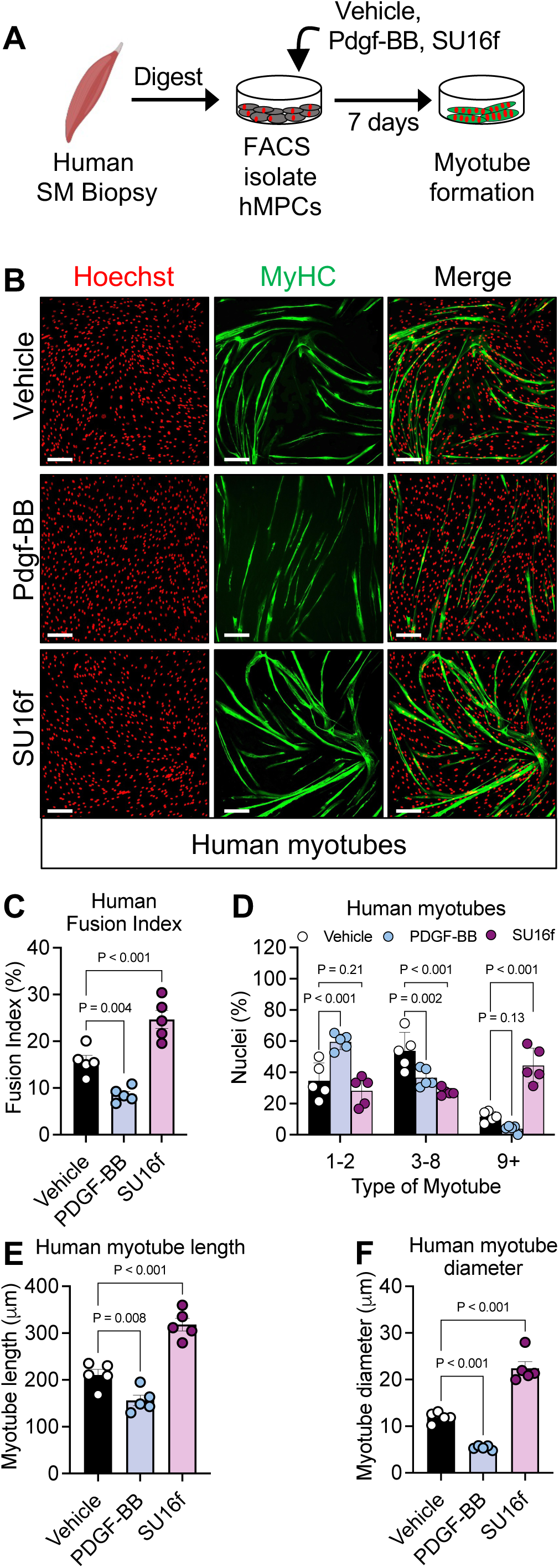
Blocking Pdgfrβ signaling in human muscle cells boosts myotube development. (A) Schematic of the experimental approach. Muscle progenitor cells were FACS-isolated from human quadricep muscles and cultured. These cells were treated with vehicle, PDGF-BB (25 ng/mL), or SU16f (1 µM) throughout the myogenesis process. Myotube development was assessed to evaluate the effects of these treatments. (B) Representative images of myotube development as assessed by MyHC immunostaining from human muscle cell cultures described in (A). (C) Quantification of the fusion index from cultures and images described in (A and B) offering insights into human muscle cell fusion under different treatment conditions (n = 5 biologically independent samples per group). (D) Quantification of nuclei and distribution within myotubes from images described in (B), reflecting myonuclear accretion in response to targeting PDGFRβ (n = 5 biologically independent samples per group). (E) Quantification of myotube length from cultures described in (A) and (B), providing a measure of myotube development under the influence of PDGF-BB and SU16f (n = 5 biologically independent samples per group). (F) Quantification of myotube diameter from the cultures described in (A) and (B), indicating the effect of the treatments on the thickness of the developing myotubes (n = 5 biologically independent samples per group). Data are presented as mean values with individual data points ± S.E.M. Statistical significance was determined using a one-way ANOVA for panels (C-F). Scale bar = 100 µm.

## Discussion

Skeletal muscle cell fusion is critical for both muscle development and regeneration^3,5,8^. While fusogenic proteins are critical for mediating cell-to-cell fusion, the regulatory pathways preparing myocytes for this process remain less well understood. This study highlights the role of PDGFRβ signaling as a negative regulator of myocyte fusion, thereby delaying muscle regeneration. Our findings reveal that heightened PDGFRβ signaling impedes myocyte fusion by reducing cell spreading and upregulating TGFβ signaling, to slow the regenerative process. Notably, these observations resonate with human gain-of-function PDGFRβ variants, such as those found in Kosaki overgrowth syndrome and infantile myofibromas^39,40^, which underscore the importance of PDGFRβ in various pathological conditions. Moreover, PDGFRβ signaling through STAT1 has been shown to mediate several aspects of Kosaki overgrowth syndrome^38^. Similarly, we find that PDGFRβ signaling within myocytes activates STAT1 to prevent cell spreading and myocyte fusion. Thus, activating STAT1 by PDGFRβ stimulation may be a hallmark of the perturb cellular state^45,66^. PDGFRβ signaling is known to activate various kinases and transcription factors which could have critical roles of cellular and molecular outcomes, especially regarding muscle cell proliferation, differentiation, and fusion^31,32^.

PDGFRβ’s involvement in cellular proliferation and migration, particularly within embryonic development and wound healing, is well-documented^28^. Specifically in skeletal muscle, several studies have shown that adding PDGF-BB stimulates the proliferation of muscle progenitors^34^. Further, PDGF-BB treatment to *Mdx* mice, a murine model of muscle dystrophy, increased the number of satellite cells and regenerative myofibers^36^. However, these changes could be secondary to the observed decrease in fibrosis and inflammation and increased vascularization^67–69^. Interestingly, treating *Mdx* mice with imatinib, a promiscuous tyrosine kinase inhibitor that has affinity for PDGFRβ signaling, can improve muscle regeneration^70^. While these pleiotropic effects of PDGF-BB administration or PDGFRβ inhibition on muscle have been observed, it is unclear if these effects generated by PDGF-BB emanate from PDGFRβ activation^31^. Our results may suggest a different role for PDGFRβ in skeletal muscle that may diverge from its traditional growth factor receptor function. In line with this notion, we do not observe an immediate impact on PAX7 cell numbers after injury, regardless of *Pdgfrβ* deletion or activation. Additionally, we did not observe obvious changes in myocyte differentiation in response to PDGFRβ signaling. This also appeared to be the case when *Pdgfrβ* was deleted within mature myofibers. When examining myocyte PDGFRβ function, we did find changes in myocyte fusion, in vitro myotube development, myonuclear accretion, and in vivo myofiber regeneration. Although we evaluated the regenerative capacity of MyoG *Pdgfrβ* genetic models, we did not assess whether a developmental phenotype could generate changes in muscle composition, strength, and systemic metabolism. Furthermore, the exact regulation of PDGFRβ within the muscle niche and its interaction with other growth factor pathways remains an open question.

The upregulation of PDGFRβ following muscle injury indicates its role as a rapid response mechanism to tissue damage^71^. Indeed, the observation that PDGFRβ signaling inhibits myocyte fusion introduces a new regulatory mechanism that ensures controlled muscle regeneration^8^. This inhibitory effect of PDGFRβ suggests it may function as a gatekeeper, preventing premature or excessive fusion that could lead to disorganized muscle tissue repair and loss of stem cell replenishment. These data suggest that PDGFRβ signaling pathway may acts as a central hub, integrating various extracellular cues necessary for the initiation of muscle regeneration. By phosphorylating STAT1, which mediates diverse cellular processes, including inflammation and immune responses, PDGFRβ signaling might prepare the myocyte for changes in the local tissue environment^72^. These niche-derived signals from immune cells and fibroblasts could serve as centers for PDGFRβ ligands (activation or inhibition)^73,74^. The exact regulation of PDGFRβ within the muscle niche and its interaction with other growth factor pathways remains an open question. Additionally, this balance of PDGFRβ signaling is likely context-dependent, with potential different roles at various stages of regeneration and within distinct cellular environments^75^. Indeed, while our studies reveal signaling crosstalk between STAT1 and TGFβ, it remains unknown whether these pathways converge on similar genetic targets or pathways. Further investigation into STAT1 regulatory genes in myocytes could help reveal additional mechanisms mediated by PDGFRβ signaling that influence myocyte fusion.

Pharmacological inhibition of PDGFRβ in murine and human models enhances myotube development and myonuclear accretion, highlighting the potential for therapeutic strategies targeting PDGFRβ to improve muscle repair. In pathological conditions where muscle regeneration is compromised, such as in muscular dystrophies or severe injuries^76,77^, fine-tuning the fusion process through PDGFRβ modulation could promote the formation of functional muscle tissue and mitigate aberrant repair. This concept paves the way for innovative interventions to optimize muscle regeneration by targeting specific phases of the regenerative process^78^. Nevertheless, our study establishes PDGFRβ as a central regulator of muscle progenitor cell function and skeletal muscle regeneration. By modulating fusion, PDGFRβ signaling ensures proper myocyte fusion and controlled myofiber repair. Moreover, our studies highlight the potential therapeutic avenue of pharmacologically blocking PDGFRβ to enhance myocyte fusion to boost muscle regeneration and repair after injury.

## Author Contributions

S.X., and A.M.B performed the experiments, collected data, and analyzed the results. N.J.K analyzed RNA sequencing results. J.B and A.T.M. collected, expanded, and prepared human muscle cells. B.D.C. provided assistance and consultation on study design and experimental methodologies. S.X., A.M.B., and D.C.B conceived and designed the study and wrote the manuscript.

## Acknowledgements

The authors thank the Berry lab members for helpful advice and discussions. We especially acknowledge all the undergraduate researchers for performing blinded image quantification (Crystal Pascual, Chris Moon, Kaitlin Chang, Alexandra Castroverde, Amber Lindsay Prasad; Alida Pahlevan Sabbagh; Jade Lindsay Palmer-Johnson). The authors thank Heather Roman for early work on establishing methodologies and animal breeding. We thank the Cornell Biotechnology Resources Center Flow Cytometric Core Facility and the Center of Animal Resources and Education for excellent assistance. We thank the Transcriptional Regulation and Expression Facility (TREx) core facility for conducting the bulk RNA sequencing. This work was supported by Cornell University institutional funds to D.C. B and D.C.B is supported by the National Institutes of Health from NIDDK [R01-DK132264].

## Materials and Methods

### Sex as a biological variable

Male and female mice were used in this study. Human muscle samples were collected from female volunteers and cultured.

### Animals

All mice were housed and experimented upon in accordance with the guidelines of Cornell University and the Institutional Animal Care and Use Committee. Pdgfrβ^fl/fl^(Stock No: 010977), Rosa26-^tdTomato^ (Stock No: 007914), Pax7-^CreERT2^ (Stock No: 017763), and Pdgfrβ^D849V^ (Stock No: 018434) mice were obtained from Jackson Laboratory (Bar Harbor, ME, USA). Myogenin-Cre mice were generously provided by Dr. Eric Olson. HSA-MCM (Stock No: 025750^56^; Jackson laboratory) mice were generously provided by Dr. Jan Lammerding. Myogenin-Cre or Pax7-^CreERT2^ mice were crossed with either *Pdgfrβ^fl/fl^* or *Pdgfrβ^D^*^849^*^V^* mice to obtain mutant strains. Control and mutant mice were then further crossed with Rosa26-^tdTomato^ to allow for lineage tracing via tdTomato expression. Mice underwent eight generations of crossing prior to experimentation. Mice were housed at room temperature (∼23 °C; ∼35% humidity) following a 14:10 hour light dark cycle. Mice were fed *ad lib* (Chow diet: Teklad LM-485 Mouse/Rat Sterilizable Diet). To induce recombination, tamoxifen (Cayman Chemical 13258) was prepared in sunflower oil (Sigma S5007) and injected intraperitonially for two consecutive days at a dose of 50 mg/kg unless otherwise noted. For pharmacological treatments, mice were intraperitoneally injected with one dose of vehicle (0.1% DMSO), PDGF-BB (50 ng/mouse), SU16f (2 mg/Kg, or fludarabine (3 mg/Kg) for five consecutive days, starting 24 hours after muscle injury. All experiments were performed on either male or female mice ranging from postnatal days 45-60.

### Human MuSC Culture

Human MuSCs were generously obtained from the laboratory of Dr. Anna Thalacker-Mercer^79^. Briefly, young (21–40 years) were recruited from the Tompkins County, New York area. Participants were excluded if they had a history of negative or allergic reactions to local anesthetic, used immunosuppressive medications, were prescribed anti-coagulation therapy, were pregnant, had a musculoskeletal disorder, suffered from alcoholism (>11 drinks per week) or other drug addictions, or were acutely ill at the time of participation. The Cornell University Institutional Review Board approved the protocol, and all the subjects provided written informed consent in accordance with the Declaration of Helsinki. Muscle biopsies were taken from the vastus lateralis muscle and cells were isolated, cultured (DMEM/F12, 20% FBS with bFGF at 5 ng/ml), and passaged four times. From these cultures, human muscle progenitor cells were FACS isolated based on CD54+ and CD29+. Cells were replated in growth media (DMEM/F12, 20% FBS and bFGF at 5 ng/ml) and cultured twice. Cells were grown to confluency on 1.4% rat collagen coated plates. Cells were then switched into differentiation media consisting of DMEM/F12 with 5% horse serum, with the media replenished daily along with denoted ligands. Myotube development was assessed seven days later^79^.

### Skeletal Muscle Injury

Mice were anesthetized by administering isoflurane at 3% for 5 minutes. The lower hindlimb was shaved and skin was sterilized with ethanol prior to injury. Injuries were inflicted at the denoted time point by directly injecting 50 µL of 1.2% Barium Chloride (BaCl_2_, Ricca Chemical, USA) into the left tibialis anterior (TA) muscle. Mice were euthanized at either 5- or 7-days post-injury (d.p.i.).

### Histological Analysis

The TA muscles were surgically removed and embedded horizontally in OCT compound (Tissue-Tek® O.C.T. Compound, Sakura Finetek, USA) using disposable embedding molds (Epredia™ Peel-A-Way™, VWR, USA), snap-frozen with liquid nitrogen, and stored at -80 °C prior to sectioning.

### Immunofluorescent Staining

Tissues were sectioned at a thickness of 20 µm using a Leica CM1950 cryostat. The sections were air-dried for 10 minutes, followed by OCT removal, and hydrophobic circles drawn around the tissues using a PAP pen (SPM0928; IHC World LLC). Sections were then fixed with 4% paraformaldehyde (PFA; 15700 Electron Microscopy Sciences) for 20 minutes at room temperature, then washed with staining buffer (0.1% Sodium Azide and 1% FBS in 1x PBS, pH 7.4) for 5 minutes. To quench the fixation, 0.1 M glycine (in 1x PBS, pH 7.4) was added for 10 minutes. Next, tissues were washed twice for 5 minutes with staining buffer and permeabilized with 0.25% Triton in staining buffer. The tissues were blocked with 5% BSA for 30 minutes at room temperature, then incubated with primary antibodies (laminin 1:200, CAT# L9393, and eMyHC 5 µg/mL, DSHB, F1.652), which were diluted in staining buffer, at 4 °C overnight. After removing the primary antibodies, tissues were washed twice for 5 minutes with 1x PBS. Secondary antibodies (anti-mouse IgG1 647, anti-rabbit 488 or 526, Invitrogen, USA) were diluted in 1x PBS at a 1:200 ratio and incubated with tissues for 2 hours at room temperature. Nuclei were stained with Hoechst (H3570; Life Technologies) at 1 µg/mL (in 1x PBS) for 10 minutes at room temperature, then washed 3 times for 5 minutes and mounted with Thermo Scientific™ Shandon™ Immu-Mount™ mounting media. Fluorescent images were taken using a Leica DMi8 inverted microscope system. Three images per biological replicate were taken for quantification, with at least three biological replicates used per experiment. For Pax7 immunostaining, cryosections were air-dried for 1 hour at room temperature, followed by 7 minutes of fixation using 4% PFA. Samples were then washed 3 times with 1x PBS for 5 minutes per wash. Tissues were blocked with 3% BSA in 1x PBS for 1 hour at room temperature. Laminin (Rabbit Ab, cat # L9393) was diluted 1:50 in blocking buffer and tissues incubated overnight at 4°C. After washing 3 times for 5 minutes each, secondary antibody (AlexFluo 488 Donkey anti-rabbit, Invitrogen) was diluted 1:250 in 1x PBS and incubated with tissues for 1 hour at room temperature. Tissues were washed twice for 5 minutes with 1x PBS, followed by a 5-minute wash with 1x R-Buffer A (Electron Microscopy Sciences, USA). Antigen retrieval was performed with 1x R-Buffer A at 95°C for 11 minutes. Slides were cooled to room temperature, washed 3 times for 5 minutes each with 1x PBS, and blocked for 1 hour in M.O.M Ms IgG blocking buffer (1 drop/ml 1x PBS) at room temperature. Tissues were then blocked for 1 hour in 10% goat serum in 1x PBS at room temperature. PAX7 antibody (2 µg/ml in 10% goat serum, DSHB) was incubated for 1 hour at room temperature and then transferred to 4°C overnight. Slides were washed 4 times for 5 minutes each with 1x PBS, incubated for 70 minutes in goat anti-mouse biotinylated secondary antibody (1:1000) in 10% goat serum, and washed 3 times for 3 minutes each in 1x PBS. Slides were then incubated for 1 hour in 2-3 drops per slide of HRP-conjugated streptavidin (Alexa Fluor 594 Tyramide Superboost Kit, Invitrogen), washed 3 times for 10 minutes each in 1x PBS, and incubated in tyramide working solution for 8 minutes. Stop solution (100 µl) was added to the tyramide reaction buffer to terminate the reaction, followed by washing 3 times for 5 minutes each with 1x PBS. Tissues were post-fixed with 4% PFA for 5 minutes, washed 3 times for 3 minutes each with 1x PBS, and incubated with Hoechst (H3570; Life Technologies) at 1 µg/ml (in 1x PBS) for 10 minutes at room temperature. After washing 3 times for 5 minutes each in 1x PBS and once with distilled water, tissues were mounted with Thermo Scientific™ Shandon™ Immu-Mount™ mounting media. Fluorescent images were collected using a Leica DMi8 inverted microscope.

### Histological Quantification

#### PAX7 Number Quantification

For analysis of satellite cell numbers, cross-sections from the TA were stained for PAX7, Laminin, and Hoechst. All PAX7-positive nuclei were manually counted by an observer blinded to genotype and age, and the counts were divided by the total area of muscle imaged per section, using Fiji ImageJ.

#### Embryonic Myosin Heavy Chain (eMHC) Quantification

Quantification of eMHC was performed on TA muscles with 70% of total muscle injured. 3 representative images were taken from each biological replicate with at least 3 biological replicates quantified per experiment. The number of eMHC-positive fibers was counted and normalized to the total number of injured muscle fibers, which were identified by the presence of centralized nuclei. The results are expressed as the percentage of eMHC-positive fibers out of the total injured fibers, using Fiji ImageJ.

#### CSA quantification

Cross-sectional area, as delineated by laminin staining, was quantified using Fiji ImageJ by manually circling each fiber to obtain the area with each muscle fiber. For injury studies, only injured fibers with centralized nuclei were quantified; while for non-injured mice, all the imaged fibers were quantified. 3 representative images were taken from each biological replicate with at least 3 biological replicates quantified per experiment.

#### Nuclei per Injured Myofiber

Nuclei within injured myofibers were manually quantified and classified as having 1, 2, 3, or 4 centrally located nuclei, then divided by the total number of injured fibers per image. 3 representative images were taken from each biological replicate with at least 3 biological replicates quantified per experiment.

### Muscle Stem Cell Isolation

The gastrocnemius, quadricep, and hamstring muscles of the hindlimbs were dissected, weighed, minced and digested with 20 mg/mL collagenase D (Roche 11088882001) in DMEM/F12 and dispase II (Roche 04942078001) at 8 units/ml at 37°C with gentle rocking for 120 minutes total. Following digestion, samples were passed through a 70 µm cell strainer and centrifuged at 500 x g for 6 minutes. Pellets were resuspended in 1X red blood cell lysis buffer (BioLegend 420301) for 5 minutes on ice. 5 mLs of DMEM/F12 media supplemented with 10% fetal bovine serum (FBS) was added to quench the reaction, followed by passing through 40 µm cell strainer and samples were then centrifuged again at 500 x g for 6 minutes. To enrich for MuSCs, pellets were then resuspended in 1 mL of antibody cocktail consisting of biotinylated antibodies against CD45 (BioLegend 103104), CD11b (BioLegend 101204), CD31 (BioLegend 102404), and Sca1 (BioLegend 108104) and incubated for 20 minutes on ice. 5 mls of DMEM/F12 media supplemented with 10% fetal bovine serum (FBS) was added and the suspension was then centrifuged at 500 x g for 6 minutes. The pellet was then resuspended in 2 mL of MACS buffer (Hanks Buffered Salt Solution, 2% FBS, 2 mM EDTA) and incubated with streptavidin coated magnetic beads (Invitrogen MSNB-6002-74), spinning for 10 minutes at room temperature. The total suspension was then placed on a magnet for 5 minutes at room temperature, allowing for collection of unbound MuSCs. Unbound MuSCs were then collected, 5 mL of DMEM/F12 media containing 10% FBS added, and the suspension centrifuged at 500 x g for 6 minutes. The pellet was then resuspended on non-collagen coated plates to further select for MuSCs. Post 1-hour incubation at 37°C in 5% CO2, media containing MuSCs was collected for further experimentation.

### Cell Culture

#### Primary MuSC Culture

Post isolation, primary isolated MuSCs from mutant and control tdTomato+ mice were resuspended in growth media containing DMEM/F12, 10% FBS and bFGF (Peprotech 100-18B) at 5ng/mL. MuSCs were counted via a hemocytometer and plated at 50,000 cells per well on collagen (Corning 354236) coated plates. To assess differentiation, cells were instead plated at a lower density (10,000 cells per well) to prevent fusion. Cells were incubated at 37°C in 5% CO2 with the growth media changed 72 hours post plating. To induce recombination in vitro, cells were treated with 1uM TMX daily and grown to confluency. Cells were then switched into differentiation media consisting of DMEM/F12 with 5% horse serum, to induce myogenesis, with the differentiation media replenished daily. Cells were differentiated for 5 days prior to fixation, staining and imaging. For cell mixing (chimera) experiments^22^, Control^MyoG^, PdgfrB^D849V^, or wild type cells were isolated as previously described. Control^MyoG^ and PdgfrB^D849V^ cells were plated at 50,000. PdgfrB^D849V^ and wild-type cells (25,000 cells/each genotype) were combined and then differentiated.

#### C2C12 Culture

C2C12 cells were purchased from the ATCC (Catalog # CRL-1772). C2C12 cells were grown in DMEM with 10% FBS until 70% confluency on non-collagen coated plates. Cells were not used beyond five passages. To induce myogenesis, confluent C2C12s were switched into DMEM containing 2% horse serum for five days and media was replaced daily. Subsequently, C2C12 cells were fixed with 4% PFA for 45 minutes at room temperature prior to staining and imaging.

#### In Vitro Pharmacological Treatments

For the denoted experiments, cells were treated with either vehicle (0.1% DMSO), 25 ng Pdgfrβ ligand (PDGF-BB; VWR 10780-774), 1 µM SU16F (Tocris 3304), or 1 µM Fludarabine (Tocris Bioscience: 3495). For all experiments, cells were treated at the beginning of the differentiation, with differentiation media plus treatment replenished each day up until fixation.

### In Vitro Immunostaining

Post fixation, cells were immunostained for Myosin Heavy Chain (Invitrogen MYSN02) or Myogenin (14-5643-82 (F5D), eBioscience). Cells were washed 3 times for 5 minutes with 1X TBS, permeabilized for 30 minutes in 1X TBS with 0.3% triton x-100, then blocked for 30 minutes in 1X TBS with 5% donkey serum, all at room temperature. Cells were then incubated at 4°C overnight with primary antibody at 1:100 in blocking buffer The following day cells were washed 3 times for 5 minutes with 1X TBS then incubated with Cy5 Donkey Anti-Mouse (Jackson ImmunoResearch 715-175-150) at 1:100 in 1X TBS for 2 hours in the dark at room temperature. Cells were washed 3 times for 5 minutes with 1X TBS then stained with Hoechst at 1:1000 in 1X TBS for 10 minutes at room temperature. Cells were then imaged using a Leica DMi8 Inverted Microscope. For quantification, at least 2-3 20X images per biological replicate were taken. Images were quantified using Fiji ImageJ.

### In Vitro Quantification

#### Fusion Index

The fusion index was quantified as the total number of nuclei contained within MyHC+ or tdTomato+ myotubes, with 3 or more nuclei, divided by the total number of nuclei within the image. 2 or 3 20X mages per biological replicate were taken for quantification with at least 3 biological replicates per experiment. Images were quantified using Fiji ImageJ.

#### Differentiation Index

The differentiation index was quantified using low density cultures to prevent fusion events from occurring. The total number of nuclei within MyHC+ or tdTomato+, including mononucleated, cells over the total number of nuclei within the image was calculated. 2 or 3 20X mages per biological replicate were taken for quantification with at least 3 biological replicates per experiment.

#### Myotube Classification

Myotube classification was calculated as the number of MyHC+ or tdTomato+ cells having 1-2, 3-8, or 9 or more nuclei over the total number of MyHC+ or tdTomato+ within each image. 2 or 3 20X mages per biological replicate were taken for quantification with at least 3 biological replicates per experiment. Images were quantified using Fiji ImageJ

#### Myotube Diameter and Length

Only mature myotubes, those containing 3+ nuclei, were quantified for myotube diameter and length. Myotube diameter was calculated by manually measuring the diameter of the thinnest, thickest, and a representative segment of each mature myotube found within each image, then calculating the average. Myotube length was calculated by manually tracing the length of all mature myotubes found within each image, then calculating the average myotube length across the image. 2 or 3 20X mages per biological replicate were taken for quantification with at least 3 biological replicates per experiment. Images were quantified using Fiji ImageJ

### RNA Isolation and qPCR

For tissues the tibialis anterior muscle from one mouse was dissected and placed into Precellys tubes containing metal beads and 1 ml of TRIzol (ambion 15596). Tissues were then homogenized in a Precellys 24 homogenizer. For cells, TRIzol was added directly to the culture dish or pellet. RNA was extracted using standard chloroform extraction and isopropanol precipitation. RNA was then washed 2 times using 70% ethanol. The concentration and quality of RNA was determined using a TECAN infinite F-nano+ spectrophotometer. 1 µg of RNA was converted to cDNA using the high-capacity RNA to cDNA kit (Life Technologies #4368813). cDNA was diluted 1:10 and added to PowerUp™ SYBR™ Green Master Mix (Life Technologies A25742) with the specified primers. An Applied Biosystems QuantStudio™ 3 Real-Time PCR System was sed to perform qPCR analysis using the ΔΔ-CT method compared to the internal control, Rn18s. All datapoints represent a single biological replicate. All biological replicates were performed in 4 technical replicates. Primer sequences used: Rn18s: Forward “GTAACCCGTTGAACCCCATT”, reverse “CCATCCAATCGGTAGTAGCG” Pdgfrβ: Forward “AGGGGGCGTGATGACTAGG”, reverse “TTCCAGGAGTGATACCAGCTT”; Pax7: forward “TCTCCAAGATTCTGTGCCGAT”, reverse “CGGGGTTCTCTCTCTTATACTCC”

### RNA Sequencing

The RNAseq project was managed by the transcriptional Regulation and Expression (TREx) Facility at Cornell University. RNA from the TA muscle was isolated as described above and submitted to the TREx Facility for further quality control analysis-chemical purity (Nanodrop) and RNA integrity (Agilent Fragment analyzer. The NEBNextPoly(A) mRNA Magnetic Isolation Module (New England Biolabs) was used to isolate PolyA+ RNA. 750 ng of RNA per sample was used for Directional RNAseq library preparation using the NEBNext Ultra II RNA Library Prep Kit (New England Biolabs). Samples were quantified with a Qubit (dsDNA HS kit; Thermo Fisher). Lastly, libraries were sequenced on an Illumina instrument. 20 million reads were generated per library. The TREx Facility performed all basic analysis including preprocessing the data, mapping to reference mouse genome, and gene expression analysis to generate normalized counts and statistical analysis of differential gene expression. Further downstream analysis was performed using Rstudio software. The RNA sequencing data that supports the findings of this paper have been deposited in the Gene Expression Omnibus under-accession code <u>GSE279532</u>.

### Flow Cytometry

MuSCs were isolated as described above, pelleted, resuspended in FACS buffer (2.5% fetal bovine serum; 2 mM EDTA in 1X PBS) and filtered through a 5 ml cell-strainer capped FACS tube (BD Falcon). A BD Biosciences FACSAria Fusion was used to sort tdTomato negative or tdTomato positive cells for gene expression analysis. Alternatively, cells were fixed in 4% paraformaldehyde for 30 minutes at room temperature. Cells were then permeabilized in 0.3% TritonX-100 in 1X TBS for 30 minutes at room temperature. Cells were then incubated in blocking buffer (1X PBS, 3% BSA, 5% goat serum) for 1 hour at room temperature. Blocking buffer was removed and cells were then incubated in non-conjugated Pax7 primary antibody in blocking buffer overnight at 4 C at the following dilution: 1:100 (DSHB-Pax7). The following morning, cells were pelleted and resuspended in secondary antibody (1:100 goat anti-mouse 488, Invitrogen A-11001) for 2 hours at room temperature. For conjugated antibodies, cells were incubated in antibody diluted in blocking buffer for 30 minutes on ice. The following conjugated antibodies at the denoted dilutions were used in this study: 1:200 CD140b/Pdgfrβ 488 (Invitrogen 53-1402-82). Cells were then analyzed on a Thermo-Fisher Attune NxT cytometry.

### Immunoblotting

Primary MuSCs were isolated into 10 cm plates as described above. Cells were grown to 50% of confluency then switched into differentiation media for 24 hours, followed by 6 hours of serum-free starvation. Myocytes were treated with either a vehicle of 0.1% DMSO, or 15 ng Pdgf-BB for 15 min. For Fludarabine treatment, cells were treated with Fludarabine (5 uM) of 0.1% DMSO for 24 h, followed by 12 h of serum free starvation with same treatments. Cells receive vehicle or fludarabine treatments were stimulated by either 15 ng PDGF-BB or vehicle for 15 min. Samples were then collected and lysed on ice using 200 µL RIPA lysis buffer supplemented with protease inhibitor and phosphatase inhibitor. Lysed samples were then incubated on ice for 30 minutes, spun down at 125,00 x g for 15 min at 4 °C, and the supernatant collected. Protein concentration was determined using the Pierce protein assay kit (Pierce™ BCA Protein Assay; ThermoScientific) and a TECAN infinite F-nano+ spectrophotometer to read absorbance. 50 ug of protein per sample was mixed at a 1:5 ratio of lysate and 6X SDS/DTT, then heated at 100 °C for 10 minutes. Samples and ladder (BioRad 161-0374) were loaded into a 10% separating and stacking gel and placed into a Mini-PROTEAN Tetra Electrophoresis Cell chamber (BioRad 1658004) suspended in 1x running buffer (Biorad 1610744). Samples were run for ∼2 hours at room temperature at 90 V. Protein was then transferred for 1 hour at 100 V on ice in 1X transfer buffer (BioRad 1610771) to an immobilon PSQ PVDF membrane (Millipore ISEQ0005). Membranes were then blocked for 1 hour at room temperature in 5% BSA in 1X TBS-T. Blots were then cut according to the protein of interest’s molecular weight and incubated at 4 °C overnight in their respective primary antibodies. Primary antibodies were diluted in 5% BSA in 1X TBS-T. The following primary antibodies were used in this study: rabbit anti-phosphorylated Pdgfrβ (Y1009) (1:1000; Cell Signaling: 3124S), rabbit anti-GAPDH (1:1000; Cell Signaling: 2118), and mouse anti-phosphorylated Stat1 (Tyr701) (1:1000; Invitrogen 33-3400). The membranes were then washed and incubated in secondary antibody (Anti-mouse or Anti-rabbit IgG, HRP-linked Antibody, Cell Signaling 7076S or 7074S) for 2 hours at room temperature. Membranes were then washed and submerged in a 1:1 solution of SuperSignal™ West Pico PLUS Chemiluminescent Substrate (ThermoScientific: 34580) for 3 minutes. A FlourChem E system (biotechne® proteinsimple) developer was used for image acquisition.

### Statistical Analysis

Excel was used for raw data collection, analysis, and quantification. All graphs and statical analysis were performed using GraphPad Prism 7-9 software. Unpaired two-tailed Student’s t-test and one-way ANOVA was used to determine statistical significance for group comparisons and two-way ANOVA was used for multiple group comparisons. Individual data points (sample size) are presented and plotted as means. Error bars are expressed as ± SEM. P < 0.05 was considered significant in all the experiments. The statistical parameters and the number of biological replicates used per experiment are found within the figure legends. All experiments were repeated twice with a minimum of 3 biological replicates per group per experiment. Researchers were blinded to the experimental groups during image analysis, measurements, and quantification. The Leica Application Suite X Microscope software was used for image acquisition and analyses. The flow cytometric analysis software, FlowJo version 10.8.1, BD FACSDiva Software version 9.4, and Attune Cytometric Software 5.3.2415.0 was used to analyze cell populations and antibody staining. RStudio was used for RNAseq statistical analysis, pathway analysis, and graph generation. Figures and experimental designs were developed and created in PowerPoint, using images from https://www.biorender.com/.

## Data availability

The data that support the findings of this study are available from the corresponding author (DCB) upon reasonable request.

