## Supplementary Figures for "Suppressing PDGFRβ Signaling Enhances Myocyte Fusion to Promote Skeletal Muscle Regeneration"

Supplemental Figure 1

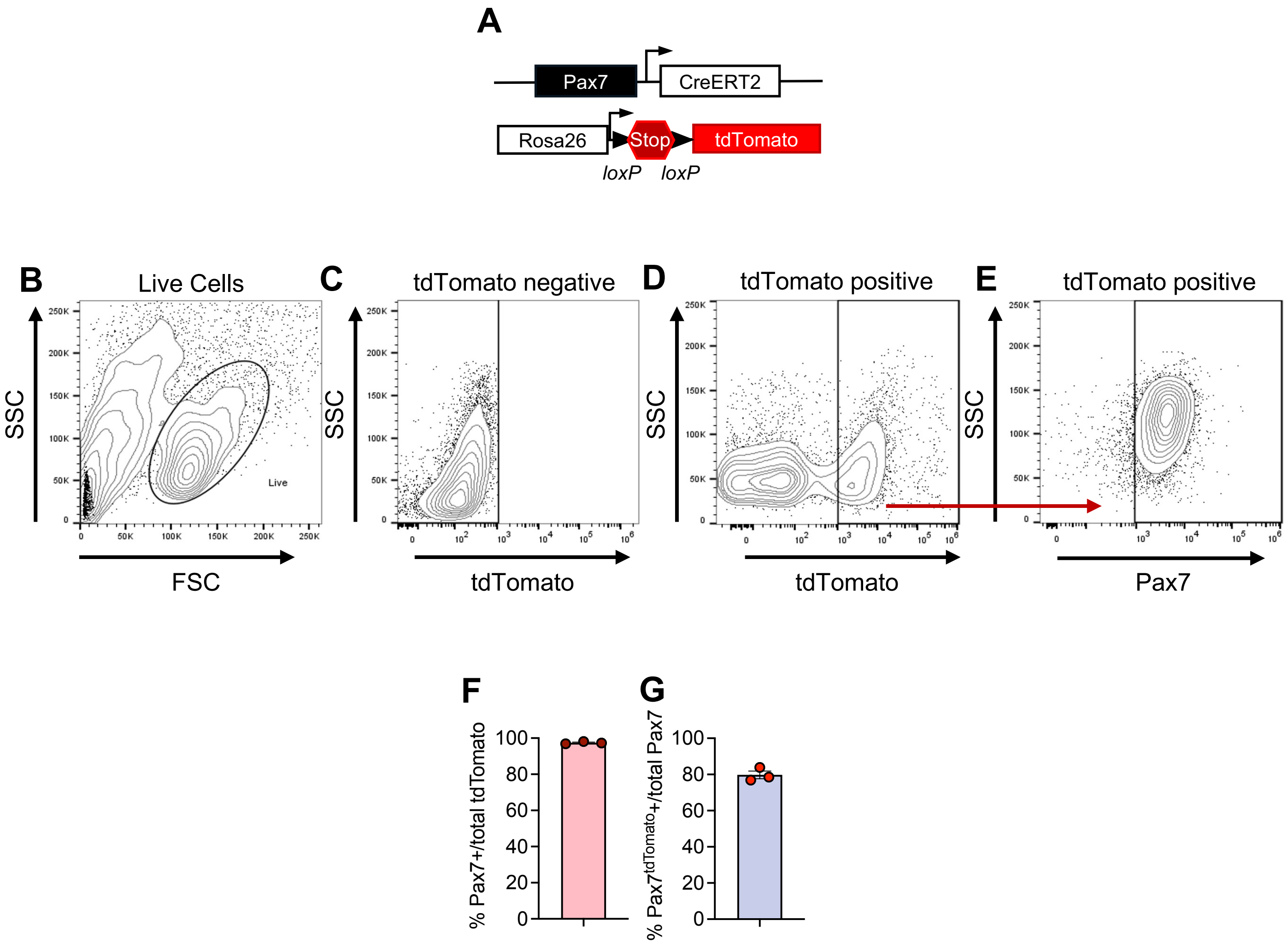

**Supplementary Figure S1. Validation of the Pax7$^{tdTomato}$ genetic tool**

(A) Allelic combination to generate the TMX inducible Pax7$^{tdTomato}$ mice.

(B) Representative FACS gating strategy for live cells.

(C) Representative FACS gating strategy for tdTomato negative cells.

(D) Representative FACS gating strategy for tdTomato positive cells.

(E) Representative FACS plots demonstrating that tdTomato positive cells are PAX7 antibody positive.

(F) FACS quantification of tdTomato cells expressing *Pax7*, demonstrating reporter overlapping with endogenous PAX7 expression (n = 3 biologically independent mice/group).

(G) FACS quantification of tdTomato cells out of total PAX7 antibody expressing cells, demonstrating high recombination efficiency (n = 3 biologically independent mice/group).

Data are presented as mean ± SEM, with each experimental group comprising biological replicates as indicated.

Supplemental Figure 2

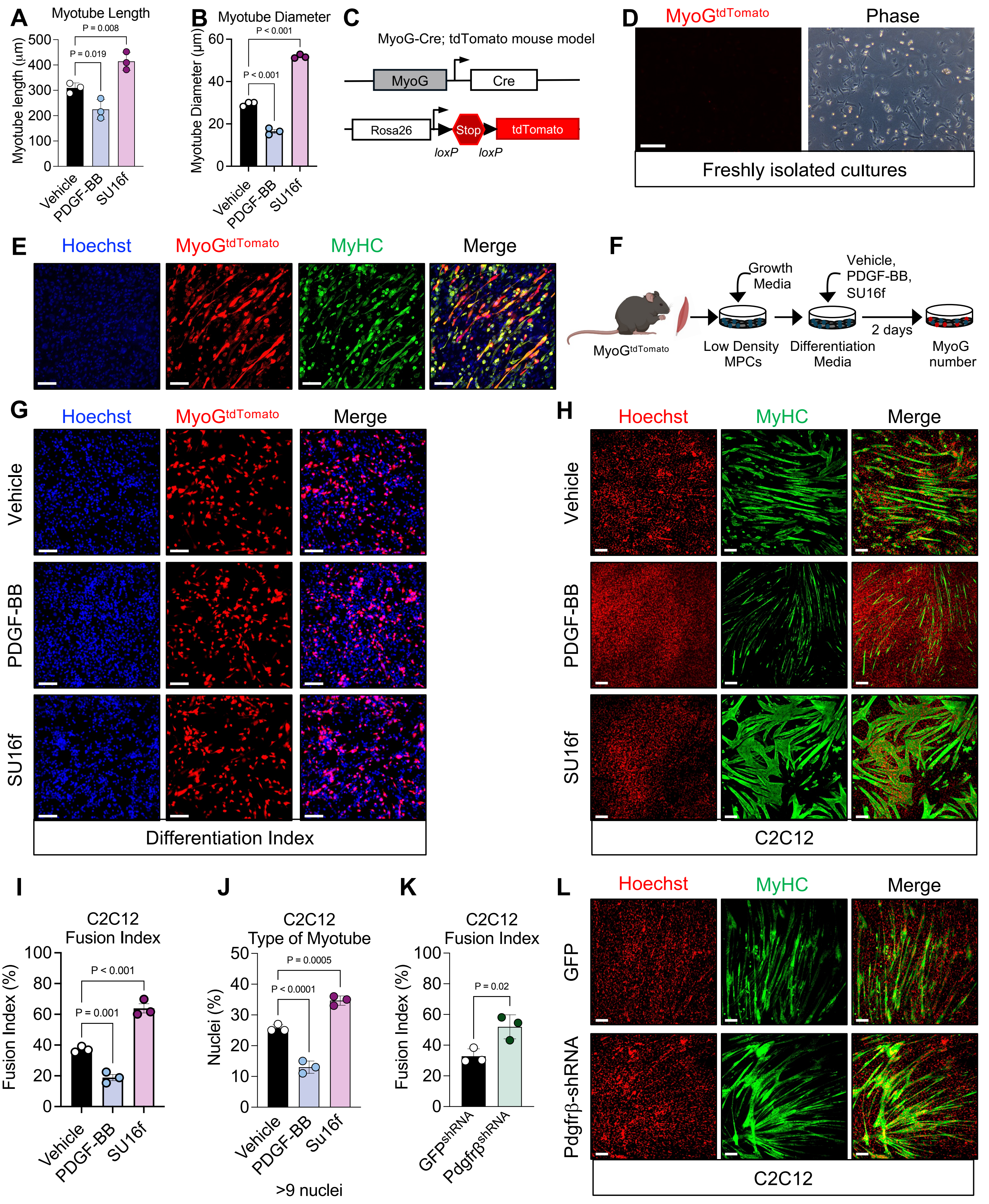

**Supplementary Figure S2. PDGFRβ activation alters myotube development**

(A) Quantification of myotube length from cultures treated with vehicle, PDGF-BB, and SU16f (n = 3 biologically independent samples per group).

(B) Quantification of myotube diameter from cultures treated with vehicle, PDGF-BB, and SU16f (n = 3 biologically independent samples per group).

(C) Allelic combination to generate the $MyoG^{tdTomato}$ mice.

(D) Muscle progenitor cells were isolated from the hindlimb muscle groups of from $Control^{MyoG}$ and visualized for tdTomato fluorescence 72 hours after plating.

(E) Representative images of myotube cultures from $Control^{MyoG}$ mice. Cultures were visualized for tdTomato fluorescence overlap with MyHC immunostaining. Note the high correspondence between tdTomato and MyHC.

(F) Schematic of the in vitro experimental procedure. Muscle progenitor cells were isolated from the hindlimb muscle groups of from $Control^{MyoG}$. Cells were plated at low density, grown, and differentiated in the presence of either vehicle (1% DMSO), PDGF-BB (25 ng/mL), or SU16f (1 µM). Myocyte formation was assessed by the number of $MyoG^{tdTomato}$ positive cells.

(G) Representative images of $MyoG^{tdTomato}$ from myocyte differentiation cultures described in (F), evaluating the role of PDGF-BB and SU16f on myocyte differentiation.

(H) Representative of images of C2C12 myotube development in the presence of vehicle (1% DMSO), PDGF-BB (25 ng/mL), or SU16f (1 µM). C2C12 derived myotubes were immunostained for MyHC to evaluate myotube development.

(I) Quantification of the fusion index from C2C12 myotube cultures described in (H), providing a measure of muscle cell fusion in response to PDGFRβ inhibition (n = 3 biologically independent mice per group).

(J) Quantification of myotubes with ≥9 nuclei per tube from cultures described in (H) (n = 3 biologically independent mice per group).

(K) Quantification of the fusion index from C2C12 myotube cultures knocked downed for *Pdgfrβ* described in (L), providing a measure of muscle cell fusion in response to *Pdgfrβ* knockdown (n = 3 biologically independent mice per group).

(L) Representative of images of C2C12 myotube development in the presence of GFP or *Pdgfrβ*-shRNA. C2C12 derived myotubes were immunostained for MyHC to evaluate myotube development.

Data are presented as mean values with individual data points ± S.E.M. Statistical significance was determined using a one-way ANOVA for panels (A), (B), (I), and (J) or unpaired Student t-test for panel (K). Scale bar = 100 µm.

Supplemental Figure 3

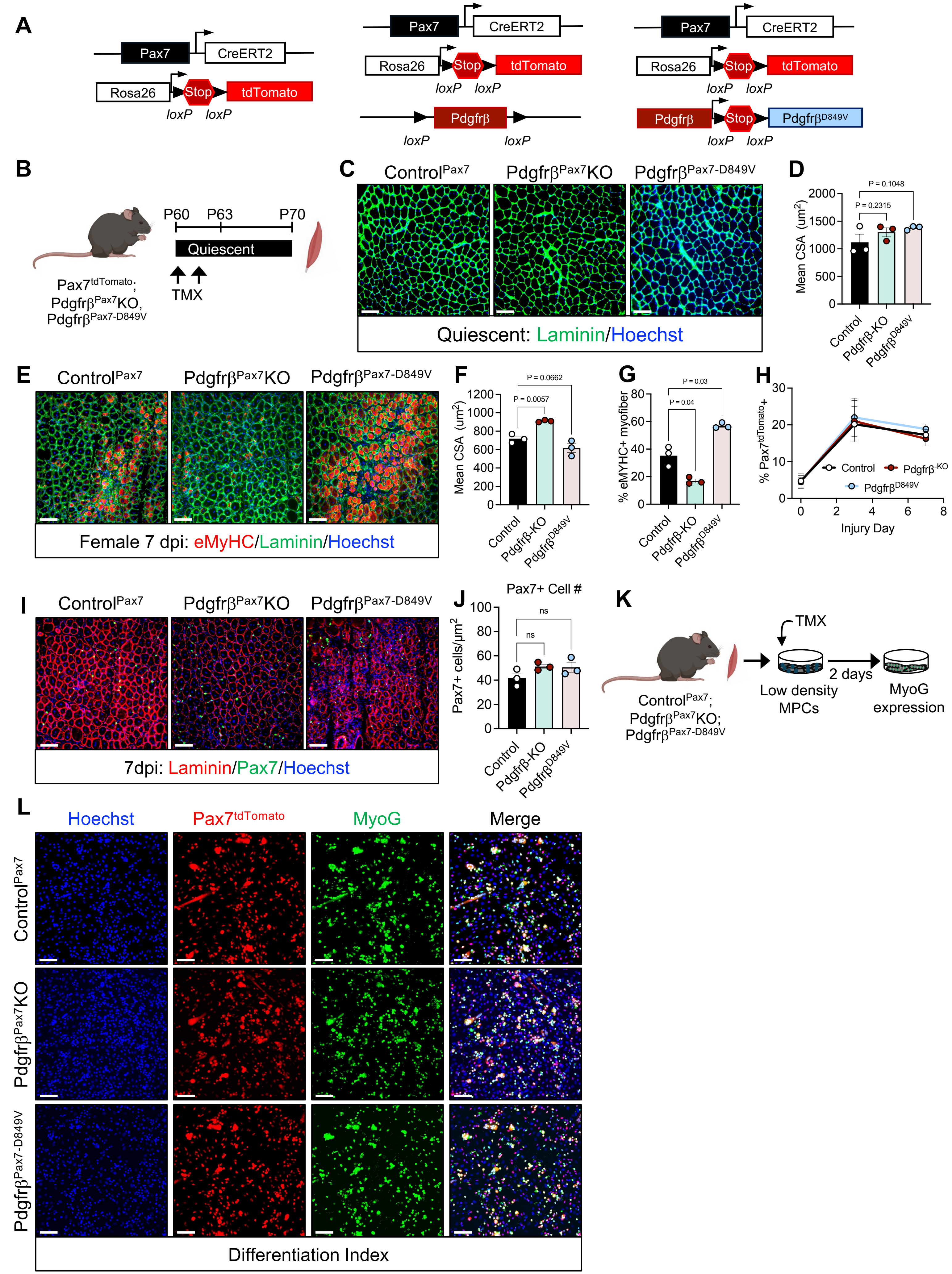

**Supplementary Figure S3. Genetically altering *Pdgfrβ* expression changes muscle regeneration and myotube development .**

(A) Schematic of allele combination to generate Control$^{Pax7}$, Pdgfrβ$^{Pax7}$KO, and Pdgfrβ$^{Pax7\text{-}D849V}$ mice

(B) Schematic of the experimental approach. Control$^{Pax7}$, Pdgfrβ$^{Pax7}$KO, and Pdgfrβ$^{Pax7\text{-}D849V}$ mice were administered TMX at postnatal day 60 (P60) to induce Cre-mediated recombination. TA muscles were harvested at P70.

(C) Representative images of quiescent TA muscle sections immunostained for laminin from mice described in (B).

(D) Quantification of mean cross-sectional area (CSA) of TA myofibers from sections described in (C) from mice described in (B) (n = 3 biologically independent mice per group).

(E) Control$^{Pax7}$, Pdgfrβ$^{Pax7}$KO, and Pdgfrβ$^{Pax7\text{-}D849V}$ female mice were administered TMX at postnatal day 60 (P60) to induce Cre-mediated recombination. TA muscles were intramuscular injected with 1.2% $BaCl_2$ to induce injury and analyzed at seven days later. Representative images of injured TA muscle sections immunostained for eMyHC and laminin from mice described in (A), illustrating the regenerative response is not sex-dependent.

(F) Quantification of mean CSA of TA myofibers from female mice and sections described in (E) (n = 3 biologically independent mice per group).

(G) Quantification of eMyHC immunostaining from TA muscle sections described in (E), indicating the level of ongoing regeneration post-injury (n = 3 biologically independent mice per group).

(H) Control$^{Pax7}$, Pdgfrβ$^{Pax7}$KO, and Pdgfrβ$^{Pax7\text{-}D849V}$ male mice were administered TMX at postnatal day 60 (P60) to induce Cre-mediated recombination. TA muscles were injured with 1.2% $BaCl_2$ and Pax7$^{tdTomato}$+ cell were FACS at zero, three, and seven d.p.i. and total Pax7$^{tdTomato}$ cell number was evaluated (n = 5 biologically independent mice/group).

(I) Representative images of injured TA muscle sections at seven d.p.i. immunostained for PAX7 from mice described in (H).

(J) Quantification of PAX7 immunostaining from TA muscle sections described in (I), indicating the number of PAX7+ cells within ongoing regeneration muscle across different genotypes (n = 3 biologically independent mice per group).

(K) Schematic of the experimental approach. Muscle progenitors were isolated from hindlimb muscle groups from of Control$^{Pax7}$, Pdgfrβ$^{Pax7}$KO, and Pdgfrβ$^{Pax7\text{-}D849V}$. After isolation, low density cultures were administered TMX to induce recombination, expanded, and subsequently differentiated and differentiation index was assessed.

(L) Representative images of colocalization of tdTomato fluorescence and myogenin-positive cells, used to calculate the differentiation index from mice described in (K).

Data are presented as mean values with individual data points ± S.E.M. Statistical significance was determined using a one-way ANOVA for panels (D), (G), and (J). Scale bar = 100 μm.

Supplemental Figure 4

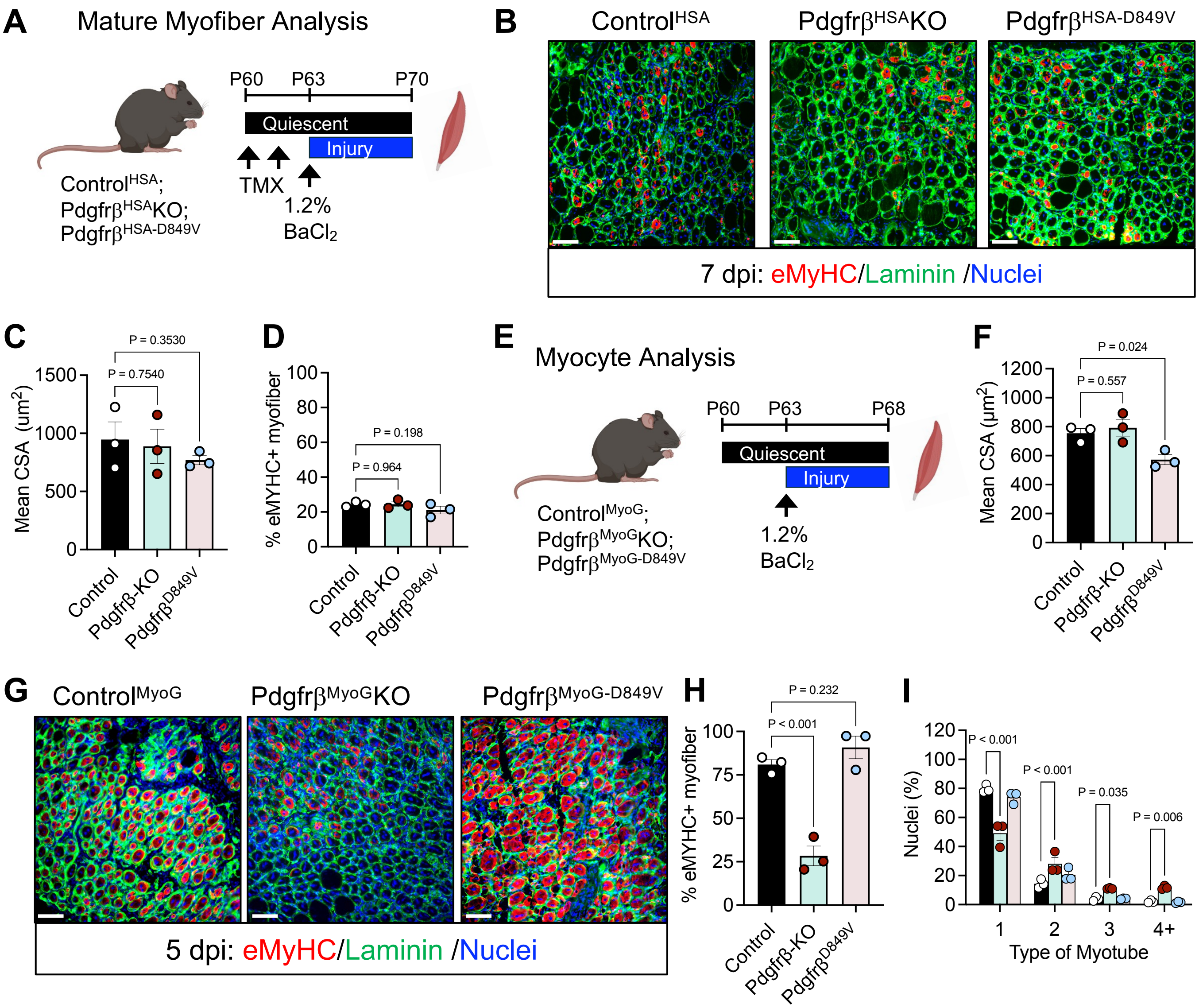

**Supplementary Figure S4. PDGFRβ regulates myocyte fusion.**

(A) Schematic of the experimental approach to evaluate Pdgfrβ function in mature myofibers. $Pdgfr\beta^{fl/fl}$ and $Pdgfr\beta^{D849V}$ alleles were combined with the HSA-MerCreMer mouse model. $Control^{HSA}$, $Pdgfr\beta^{HSA}KO$, and $Pdgfr\beta^{HSA\text{-}D849V}$ mice were administered TMX at postnatal day 60 (P60) to induce Cre-mediated recombination. TA muscles were injured with 1.2% $BaCl_2$ and analyzed at seven days later.

(B) Representative images of injured TA muscle sections immunostained for eMyHC and laminin from mice described in (A), illustrating the regenerative response across different genotypes.

(C) Quantification of mean cross-sectional area (CSA) of injured TA myofibers from sections described in (B) from mice described in (A) (n = 3 biologically independent mice per group).

(D) Quantification of eMyHC immunostaining from TA muscle sections described in (B) (n = 3 biologically independent mice per group).

(E) Schematic overview of the experimental setup: TA muscles from $Control^{MyoG}$, $Pdgfr\beta^{MyoG}KO$, and $Pdgfr\beta^{MyoG\text{-}D849V}$ mice were subjected to chemical injury using 1.2% $BaCl_2$. Muscle tissues were harvested and analyzed at five days later.

(F) Quantification of mean cross-sectional area (CSA) of injured TA myofibers from sections described in (G) from mice described in (E) (n = 3 biologically independent mice per group).

(G) Representative images of injured TA muscle sections immunostained for eMyHC and laminin from mice described in (E), illustrating the regenerative response across different genotypes at five days post injury.

(H) Quantification of eMyHC-positive fibers within the injured TA muscle from sections described in (G) from mice described in (E) (n = 3 biologically independent mice per group).

(I) Quantification of myonuclear accretion within injured myofibers from TA muscle sections described in (G) from mice described in (E), providing insights into cellular fusion and muscle repair dynamics in the different experimental groups (n = 3 biologically independent mice per group).

Data are presented as mean values with individual data points ± S.E.M. Statistical significance was determined using a one-way ANOVA for panels (C), (D), (F), (H) and (I). Scale bar = 100 μm.

Supplemental Figure 5

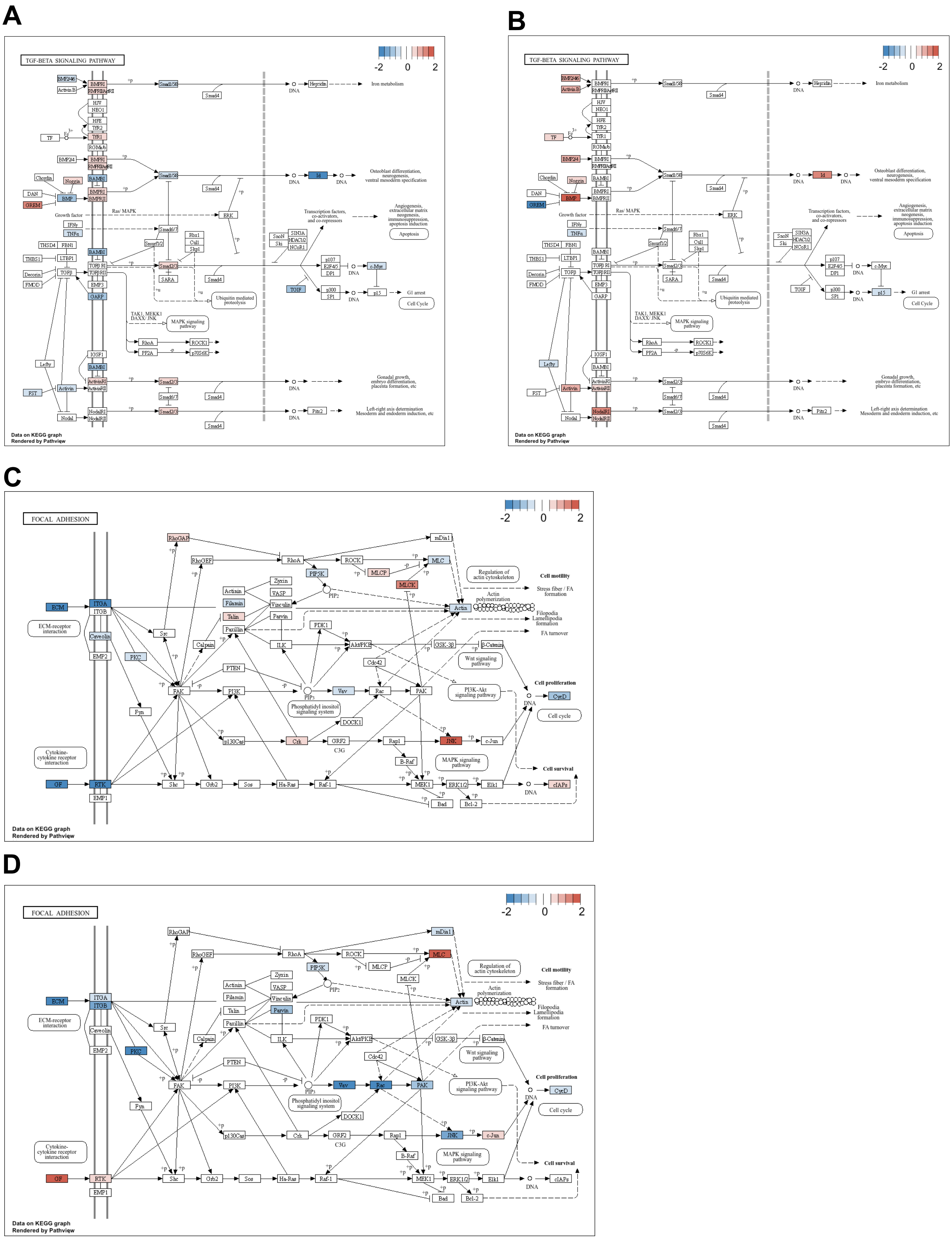

**Supplementary Figure S5. PDGFRβ cooperates with TGFβ and focal adhesion signaling pathway**

(A, B) Effected genes involved in the TGFβ signaling pathway by deleting (A) or activating (B) *Pdgfrβ*.

(C, D) Effected genes involved in the focal adhesion signaling pathway by deleting (C) or activating (D*) Pdgfrβ*.

Supplemental Figure 6

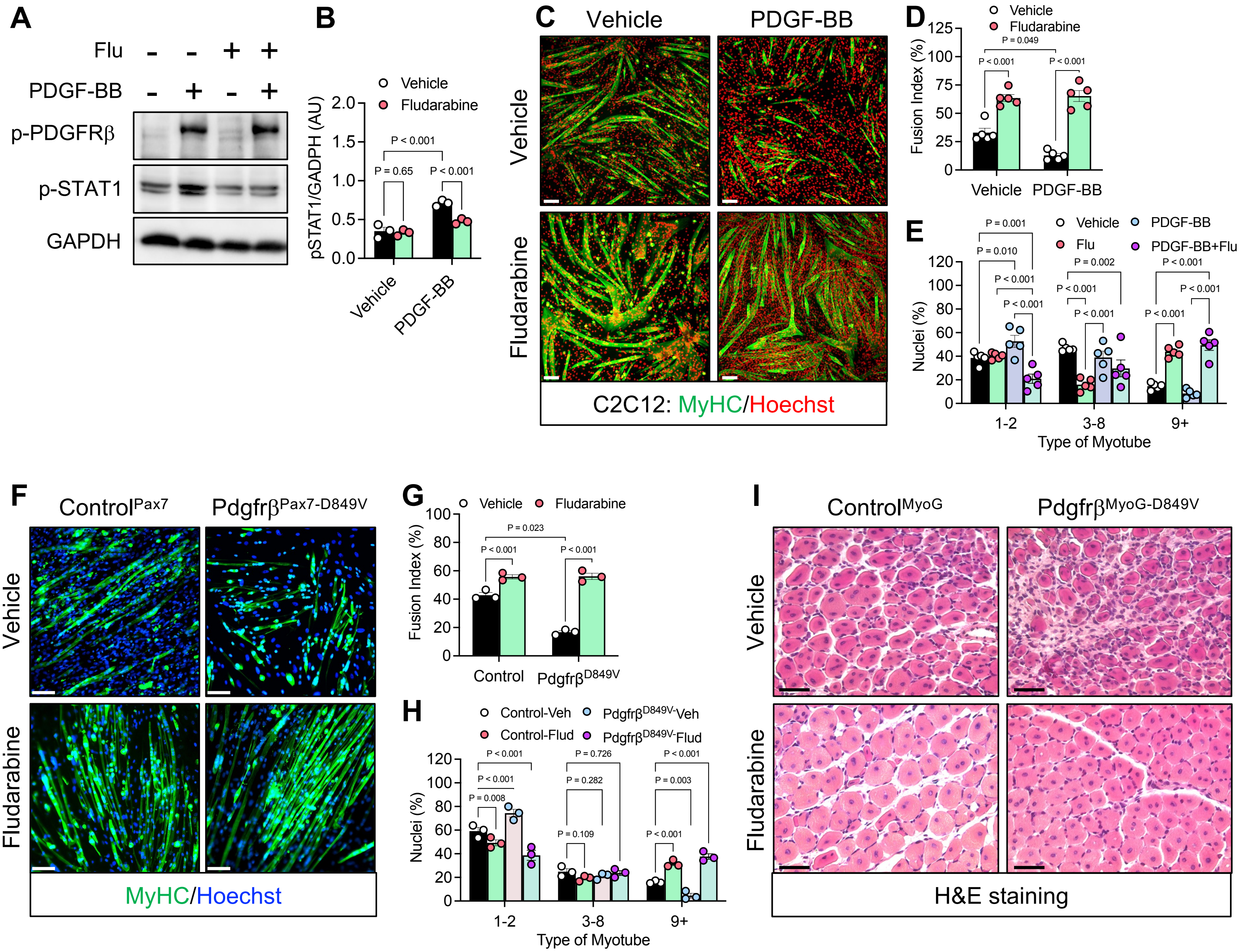

**Supplementary Figure S6. STAT1 mediates PDGFRβ signaling to mediate myocyte fusion**

(A) Muscle progenitor cells were isolated from the hindlimb muscle groups of $Control^{Pax7}$ male mice. After isolation, cells were expanded, and subsequently differentiated for one day. Myocytes were treated with vehicle, PDGF-BB (15 ng/mL), fludarabine (5 μM), or the combination of PDGF-BB and fludarabine for 15 minutes. Representative immunoblot of phosphorylated PDGFRβ and STAT1.

(B) Quantification of independent immunoblots described in (A) (n = 3 biologically independent mice/group).

(C) Representative of images of C2C12 myotube development in the presence of vehicle (1% DMSO), PDGF-BB (25 ng/mL), fludarabine (1 μM) or the combination of PDGF-BB and fludarabine. C2C12 derived myotubes were immunostained for MyHC to evaluate myotube development.

(D) Quantification of the fusion index from C2C12 myotube cultures described in (C), providing a measure of muscle cell fusion in response to STAT1 inhibition (n = 5 biologically independent mice per group).

(E) Quantification of myotube nuclei and distribution per tube type from C2C12 cultures described in (C) (n = 5 biologically independent mice per group).

(F) Muscle progenitor cells were isolated from the hindlimb muscle groups of $Control^{Pax7}$ and $Pdgfr\beta^{Pax7\text{-}D849V}$ male mice. After isolation, cells were administered TMX to induce recombination, expanded, and differentiated in the presence of vehicle or fludarabine (1 μM) for five days. Subsequently, myotube development was assessed.

(G) Quantification of the fusion index from the images described in (F), reflecting the efficiency of muscle cell fusion into myotubes (n = 3 biologically independent mice per group).

(H) Quantification of myotube nuclei count and distribution from the images described in (F) (n = 3 biologically independent mice per group).

(I) Representative images of hematoxylin and eosin (H&E) staining of injured TA muscle sections from $Control^{MyoG}$ and $Pdgfr\beta^{MyoG\text{-}D849V}$ treated with vehicle or fludarabine for five days.

Data are presented as mean values with individual data points ± S.E.M. Statistical significance was determined using a one-way ANOVA for panels (A), (B), (I), and (J) or unpaired Student t-test for panel (K). Scale bar = 100 μm.

Supplemental Figure 7

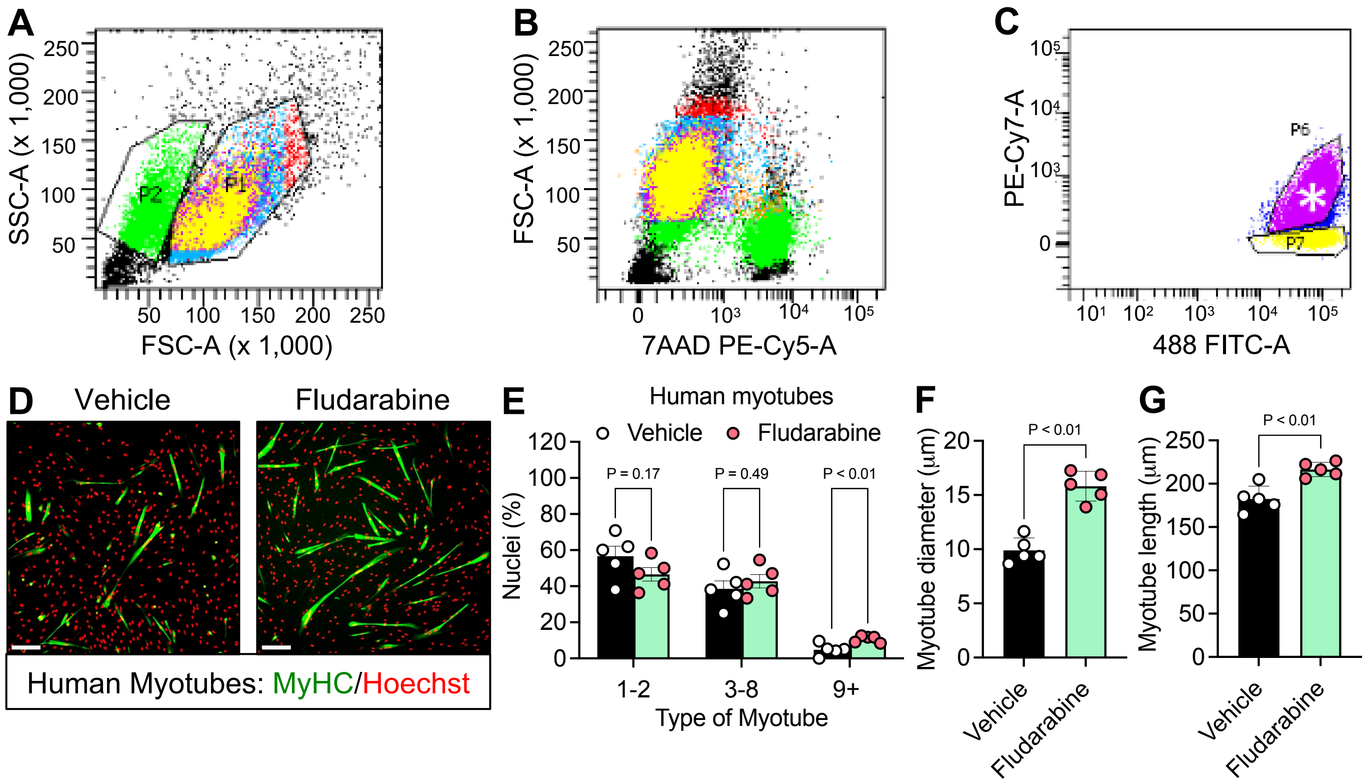

**Supplementary Figure S7. Modulating PDGFRβ signaling regulates human myocyte fusion**

(A) Representative flow cytometric plots of side and forward scatter area on live cells from human muscle samples.
(B) Representative flow cytometric plot of viability staining from isolated human muscle cells.
(C) Representative flow cytometric plots on positive selection markers CD56 and CD29.
(D) Representative images of human myotube development in cultures treated with vehicle or fludarabine (1 μM).
(E) Quantification of myotube nuclei and distribution from cultures described in (D) ( n = 5 biologically independent humans/group).
(F) Quantification of myotube diameter from cultures described in (D) (n = 5 biologically independent humans/group).
(G) Quantification of myotube length from cultures described in (D) (n = 5 biologically independent humans/group).

Data are presented as mean values with individual data points ± S.E.M. Statistical significance was determined using an unpaired Students t-test for panels (E-G). Scale bar = 100 μm.
